# Structured Schemas for Provenance-Rich, LLM-Assisted QSP Model Calibration

**DOI:** 10.64898/2026.03.05.709623

**Authors:** Joel Eliason, Aleksander S. Popel

## Abstract

Quantitative systems pharmacology (QSP) models require calibration data from literature, yet manual curation is inconsistently documented and large language model (LLM) extraction can hallucinate values and fabricate citations. We present MAPLE (Model-Aware Parameterization from Literature Evidence), which uses structured validation schemas as a collaboration interface between LLMs and modelers. Two schemas span two scales: the SubmodelTarget schema for isolated experiments constraining individual parameters, and the CalibrationTarget schema for clinical and in vivo endpoints constraining the full model. Both separate data extraction from modeling decisions, recording every value with full provenance. Targeted validators catch characteristic LLM errors by matching values to source snippets, resolving DOIs, and executing code. For a pancreatic ductal adenocarcinoma QSP model, we used MAPLE to extract and curate 37 SubmodelTargets and 45 CalibrationTargets. Before any human review, the validators triggered 50 automated retries; every value carries a direct quote from its source and a verified citation; and 11 of 19 parameters are supported by more than one independent source.The LLM drafted usable forward models and code from context, while the modeler supplied the context and scientific judgment it cannot infer, revising forward-model choices in 65% of SubmodelTargets, priors in 46%, and source relevance in all files. This evaluation covers one model in one disease area, by a single group, so it characterizes the framework rather than establishing how broadly it generalizes. MAPLE records the modeler’s reasoning in a form that can be re-run and independently checked, so it is not lost when the modeling team changes.

## 1 Introduction

Quantitative systems pharmacology models translate mechanistic understanding of disease biology into predictions of drug response ^1^. QSP models can comprise tens to hundreds of ordinary differential equations, and calibration requires data from diverse sources including targeted experimental studies and clinical endpoints ^2,3^. Because virtual population generation and virtual clinical trials depend on parameter distributions rather than point estimates ^4,5^, calibration must quantify uncertainty. As QSP increasingly integrates with machine learning ^6,7,8,9,10^, the bottleneck of manual parameter extraction becomes more acute.

Extracting calibration data from literature requires judgment about applicability, uncertainty, and potential biases, with full provenance for reproducibility. A structured representation of such data captures what was measured, under what conditions, with what uncertainty, and how that measurement relates to model parameters or observables. Some studies isolate specific mechanisms and directly constrain individual parameters, while clinical and in vivo endpoints constrain the integrated model through observables computed over the full system state.

Manual curation remains the standard approach but is labor-intensive, with inconsistent documentation across individuals and projects. When personnel change, the reasoning behind parameter choices may be lost. Without structured representations, combining constraints from multiple experiments requires ad-hoc scripting with no standard path from extracted data to inference code. Traditional NLP pipelines for automated extraction require task-specific training data ^11,12,13^ and do not generalize to the diverse parameter types in QSP modeling.

Large language models offer a more flexible approach. LLMs can identify relevant literature given mechanistic context, read scientific text, and extract information into structured templates ^14,15,16,17,18^. However, LLMs exhibit hallucination and fabrication errors that are unacceptable for quantitative modeling ^19,20,21^. Many of these failure modes, however, are amenable to systematic detection: hallucinated values can be caught by requiring verbatim source text, fabricated citations through DOI resolution, and unit and code errors by parsing units and executing the code with mock inputs. This suggests a workflow where LLMs handle extraction while automated validators catch characteristic errors, with Pydantic ^22^, a Python data validation library, enabling a self-correcting retry loop.

We present MAPLE (Model-Aware Parameterization from Literature Evidence), a framework for LLM-assisted QSP model calibration. Its core components are: (1) model-aware literature search using mechanistic context to guide LLM web search; (2) validated YAML schemas for two calibration regimes (the *SubmodelTarget* schema for isolated experiments constraining individual parameters, and the *CalibrationTarget* schema for clinical/in vivo endpoints constraining the full model), separating data extraction from model specification while preserving traceability; and (3) targeted validators for hallucination detection (value-in-snippet matching), citation verification (DOI resolution), and code correctness. The version of MAPLE used here (v0.1.0) additionally generates a joint Julia inference script from the validated targets; this inference layer has since moved to a separate package, but the analysis reported below uses the integrated v0.1.0 pipeline. The schemas, validators, and reference database do not require an LLM: a modeler can populate them manually and the validators run against the result the same way. We evaluate MAPLE on SubmodelTargets and CalibrationTargets for a pancreatic ductal adenocarcinoma (PDAC) QSP model and report metrics on validation performance and extraction reliability. The LLM produces usable forward-model choices and observable code when given enough context, but cannot tell when the context is incomplete, and it can miss inputs or lose track of constraints across long records. The schema and validators make explicit what context the modeler supplies (unit conventions, source-applicability decisions), and the modeler’s review of every record catches the LLM lapses the validators do not detect, so the calibration record shows both what the LLM produced and what the modeler supplied or corrected.

## 2 Methods

### 2.1 Model-Aware Literature Search

Figure 1 summarizes the full MAPLE pipeline. The user specifies which model parameters or observables need calibration, and the system builds a prompt that guides an LLM with web search capability to find relevant experimental literature. The framework supports multiple LLM providers (OpenAI, Anthropic, Google) through a common interface. In this study, we used GPT-5.1 for batch extraction and Claude Opus 4.6 for interactive extraction.

**Figure 1:**
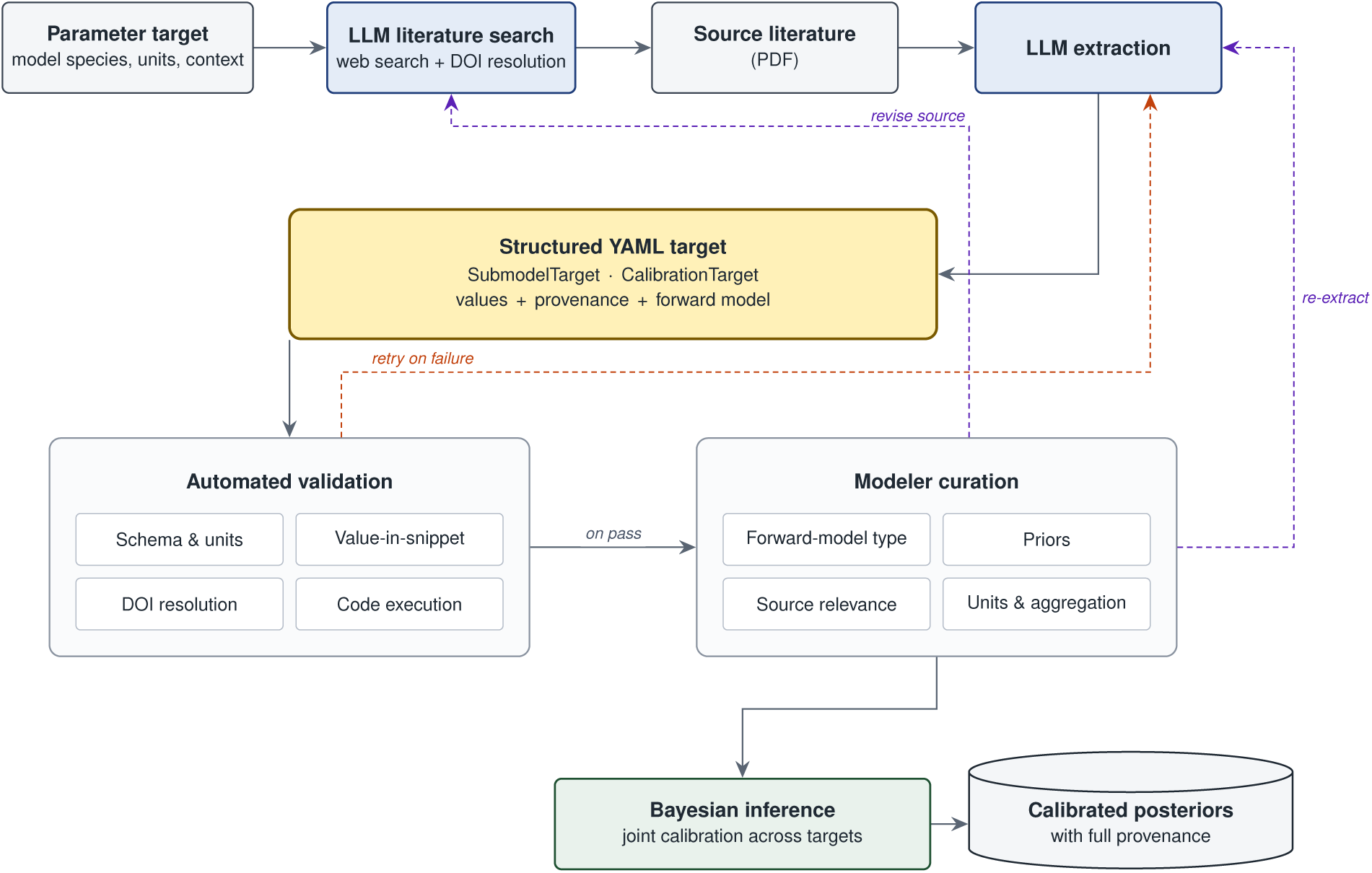
MAPLE workflow. An LLM searches the literature for a parameter target and extracts a structured YAML target carrying values, provenance, and a forward model. Automated validators gate the YAML, while failures trigger LLM retry. Validated targets are reviewed by the modeler, who may also revise the source or re-extract. Curated targets feed a joint Bayesian inference yielding posteriors with full provenance.

The prompt injects mechanistic context from the model structure: for each target, the system retrieves its name, units, and description. For parameters, it also retrieves the reactions where the parameter appears with rate laws, the species involved, and the broader reaction network. For observables, it retrieves the species and compartments that define the observable. This context guides the LLM toward experiments that constrain the targets. Source exclusion prevents reuse of papers across extractions to promote literature diversity. The prompt also includes the complete schema structure and guidance on common pitfalls, reducing the retry loop by preventing predictable errors.

### 2.2 Schema Design

The framework defines two complementary schemas for two calibration regimes. The *SubmodelTarget* schema captures isolated experiments (in vitro assays, single-mechanism studies) that constrain individual model parameters through simplified forward models. The *CalibrationTarget* schema captures clinical and in vivo endpoints that constrain the full model through observables computed over the complete species state. Both schemas share a common design philosophy but differ in how they connect literature data to model parameters and outputs. For clarity, the manuscript refers to these two schemas by name (`SubmodelTarget` and `CalibrationTarget`) rather than using “calibration target” as a general term.

#### 2.2.1 Shared Design Principles

Both schemas enforce a data-first architecture: extraction of what the paper reports is separated from specification of how to use it for calibration. Every extracted numeric value must appear verbatim in a quoted text snippet from the source, creating an auditable trail and enabling automated hallucination detection.

#### 2.2.2 SubmodelTarget: Parameter Priors from Isolated Experiments

The SubmodelTarget schema structures each record into two layers: an *inputs* layer that captures data exactly as reported in the literature (value, units, uncertainty, sample size, source reference, and quoted snippet), and a *calibration* layer that specifies how that data constrains model parameters through a simplified forward model. The submodel derives from the full QSP model by isolating relevant species. Shared parameter names establish the connection: a proliferation rate named `k_apsc_prolif` in a submodel refers to the same parameter in the full model, allowing posteriors to serve directly as priors for full-model calibration. Each parameter includes a prior distribution with documented rationale, and an error model specifies the likelihood function connecting predictions to data. Supplementary Section 6 provides detailed field descriptions with YAML examples.

##### Forward model type system

The schema provides 15 built-in forward model types, described in Supplementary Table 1. Each structured model declares its fields as typed roles rather than raw values, allowing the same template to serve different estimation scenarios depending on which field is treated as the unknown.

A complete SubmodelTarget example is provided in Supplementary Section 1; Figure 2 shows a compact version illustrating the two-layer structure.

**Figure 2:**
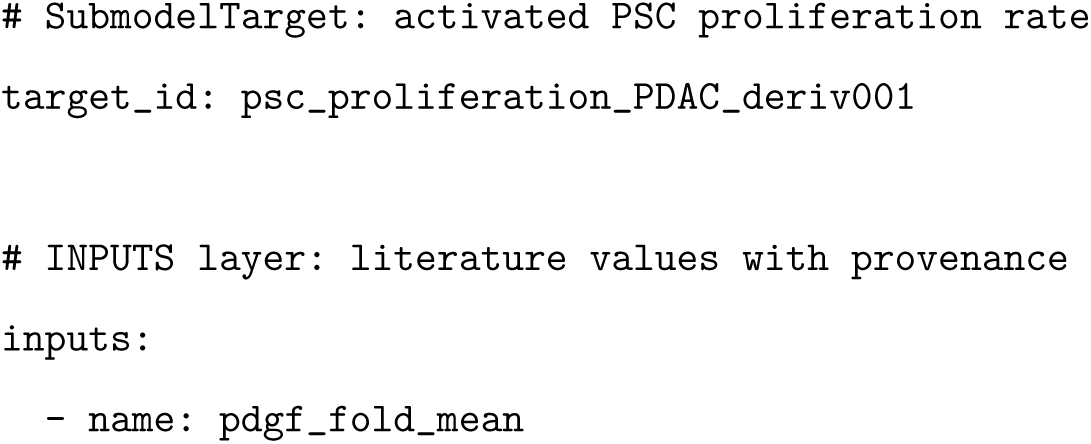
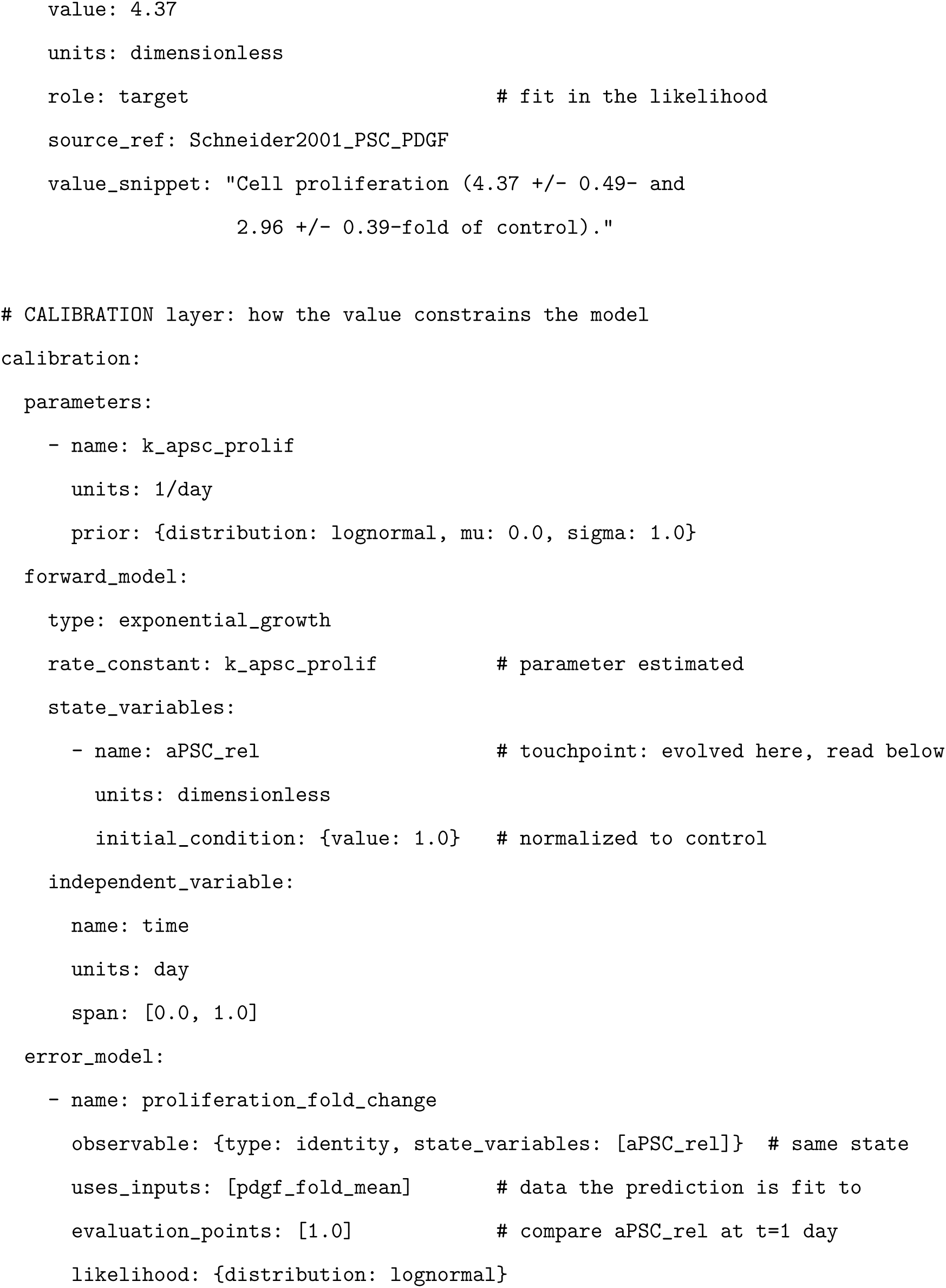
Compact SubmodelTarget example showing how an extracted value connects to the forward model. The `inputs` layer records the literature value with the verbatim snippet that anchors it. Within the `calibration` layer, the state variable `aPSC_rel` is the touchpoint between the forward model and the data: the forward model evolves it from a control-normalized initial condition at rate `k_apsc_prolif`, and the `error_model` observable reads that same state and fits its value at `t=1` day to the measured input `pdgf_fold_mean` (linked through `uses_inputs`) under a lognormal likelihood. Rationale fields and the auxiliary standard-deviation input are omitted for brevity; Supplementary Section 1 gives the complete record.

#### 2.2.3 CalibrationTarget: Clinical and In Vivo Endpoints

Many calibration constraints come from clinical or in vivo studies where the measurement reflects integrated model behavior. The CalibrationTarget schema represents these full-model endpoints through three components. The *observable* specifies how to compute the measurement from model species as a Python function that receives the solved state and returns a quantity with units; named constants carry provenance traced to a curated reference database or literature source. The *empirical data* block derives summary statistics (median and 95% CI) from literature inputs via Monte Carlo simulation. For intervention studies, a *scenario* block captures treatment context (agents, doses, schedules). Complete examples and schema definitions are provided in Supplementary Sections 7 and 5.

#### 2.2.4 Source Relevance Assessment

Experimental data rarely matches the target model perfectly. The `source_relevance` block documents the translation from source context to model context across several dimensions: indication match, species translation, evidence type, and tumor microenvironment compatibility. Each dimension requires justification when the source deviates from the target model. An `estimated_translation_uncertainty_fold` field quantifies the combined expected uncertainty from all translation factors, which propagates to prior width during inference (e.g., cross-species extraction from a proxy indication might warrant 10-fold uncertainty). Supplementary Section 13 provides detailed field descriptions.

#### 2.2.5 Schema Implementation

Both schemas are implemented using Pydantic ^22^ with typed fields and constraints validated automatically during YAML parsing. SubmodelTarget selects among its 15 forward model types with a single `type` field that determines which remaining fields are required and validated (a Pydantic discriminated union); CalibrationTarget instead has a single fixed structure, with the same source, relevance, and context fields on every target. Supplementary Section 5 provides the complete schema definitions and Supplementary Section 12 provides implementation details.

### 2.3 Validation Framework

Both schemas share a common validation core with schema-specific extensions. Each validator runs independently, and a target must pass all validators before proceeding to downstream use.

#### 2.3.1 Reference Validation

LLMs sometimes generate references to entities that do not exist elsewhere in the document. Reference validators check that internal references (`uses_inputs`, `source_ref`, model parameters, and observable state variables) all resolve to defined entities.

#### 2.3.2 Content Validation

Content validators address fabrication of values, citations, and other factual claims. *DOI validation* queries CrossRef for every DOI; title validation further checks that the extracted paper title fuzzy-matches the CrossRef metadata (75% similarity threshold), catching cases where a valid DOI is paired with fabricated bibliographic details. *Value-in-snippet validation* compares each extracted numeric value against its associated source text, providing the primary anti-hallucination defense: when an LLM hallucinates a number, it rarely fabricates a consistent snippet, and the mismatch provides a reliable detection signal. A separate external validator (Section 2.3.3) further verifies that snippets appear in the cited papers. *Unit validation* parses all unit strings using Pint ^23^ to verify dimensional validity. *Code validation* executes observation and observable code with mock data to verify correct behavior, catching logic errors and signature mismatches before inference. *Source relevance validators* enforce minimum uncertainty thresholds for cross-species (2-fold), cross-indication (3-fold), and low TME compatibility (10-fold) extractions, ensuring translation uncertainty propagates to prior widths. The fuzzy-match thresholds (75% for titles, 80% for snippets) were set to tolerate minor formatting, punctuation, and OCR differences between extracted and source text while still rejecting genuinely different strings, and are configurable defaults the modeler can tighten or loosen; the cross-species, cross-indication, and low-TME multipliers are likewise conservative minimum uncertainty defaults, applied as floors that the modeler may widen further when source applicability is weaker.

Each validator raises a specific exception class with a `category` attribute (hallucination, fabrication, code, units, reference, structural, prior) that enables aggregation across extraction runs and feeds into the metrics reported in Section 3. Optional Logfire instrumentation^24^ provides OpenTelemetry-based tracing for debugging extraction failures. Supplementary Section 3 provides implementation details for each validator.

#### 2.3.3 External Validation

An additional external validator verifies that value snippets actually appear in the cited source papers by fetching paper text from Europe PMC and Unpaywall using the DOI, then using fuzzy matching (80% similarity threshold) to locate each snippet. This catches a subtle failure mode where the LLM fabricates plausible-looking snippets that contain the correct value but do not actually appear in the cited paper. Because this requires network access to external APIs, it runs as a separate manual step. It is provided as an on-demand script (`scripts/validate_snippets_in_source.py`) in the analysis repository, built on the MAPLE full-text fetch and fuzzy-matching validators, and was applied selectively during curation rather than as part of the automated per-target validation gate, since external full-text availability is uneven across sources; the value-in-snippet check described above is the systematically applied internal anti-hallucination defense.

#### 2.3.4 Structural and CalibrationTarget-Specific Validation

Structural validators check schema-level constraints (time span ordering, required fields per model type, array dimension consistency). CalibrationTarget adds validators for full-model observables: observable code execution against mock species data, a denominator audit for density/fraction observables (catching a common source of order-of-magnitude errors), species existence checks, and distribution code verification that computed summary statistics match reported empirical data within tolerance (1% for median, 10% for CI bounds). Supplementary Section 3 provides implementation details for all validators.

### 2.4 Inference Code Generation

A validated SubmodelTarget already specifies its forward model, priors, and likelihood, so the same record that passes validation can be turned into inference code without further hand-coding. MAPLE v0.1.0 (the version used here) includes a translator that emits a Julia script for Turing.jl^25^; later versions of MAPLE keep the schema and validators and provide the inference translator in a separate package. The translator joins targets that share parameter names, so each shared parameter is declared once and constrained by every source that constrains it. Supplementary Section 4 describes the generated code structure, forward model handling, and inference diagnostics. The two schemas connect through a natural calibration pipeline: SubmodelTarget posteriors from isolated experiments become informative priors for full-model calibration against CalibrationTarget endpoints. Because SubmodelTarget parameter names match the full model, the posterior distributions transfer directly without post-hoc mapping.

### 2.5 Application: PDAC QSP Model

The PDAC QSP model used in this study is adapted from a spatial QSP model of hepatocellular carcinoma ^26^ that first incorporated fibroblast dynamics into the tumor microenvironment. We evaluate MAPLE on this model using both schemas: 37 curated SubmodelTargets for individual parameter calibration from isolated experiments (spanning tumor growth, stromal dynamics, immune recruitment, and cytokine signaling), and 45 CalibrationTargets for full-model endpoints (baseline tumor microenvironment composition, neoadjuvant immunotherapy response, and clinical progression). Batch extraction used OpenAI `gpt-5.1` at high reasoning effort, both for SubmodelTargets and for the batch-pipeline portion of CalibrationTargets; interactive CalibrationTarget extraction and all curation used Anthropic `claude-opus-4-6` through Claude Code. All extraction and curation runs were performed in January and February 2026. This study used only previously published, aggregate literature data and did not involve human subjects or animal experimentation, so institutional review board or IACUC approval was not required.

#### 2.5.1 Evaluation Metrics

We report metrics across four workflow stages: extraction performance, error analysis, source characteristics, and curation analysis.

## 3 Results

### 3.1 SubmodelTarget Extraction and Validation

We applied MAPLE to extract SubmodelTargets for a PDAC QSP model, covering tumor growth, stromal dynamics, immune recruitment, and cytokine signaling. All 37 curated SubmodelTargets used for joint inference were produced by batch extraction through the automated retry loop, followed by curation. Of these, 18 were traced with Logfire and supply the per-target retry, duration, and token metrics in Table 1; the remaining extractions predate that instrumentation and so are not in the table. The instrumented batch and the inference set therefore overlap but are not identical: a few instrumented runs targeted parameters that were consolidated or dropped during curation. The schema-validation and curation results in Section 3.2 cover all 37 targets.

**Table 1:**
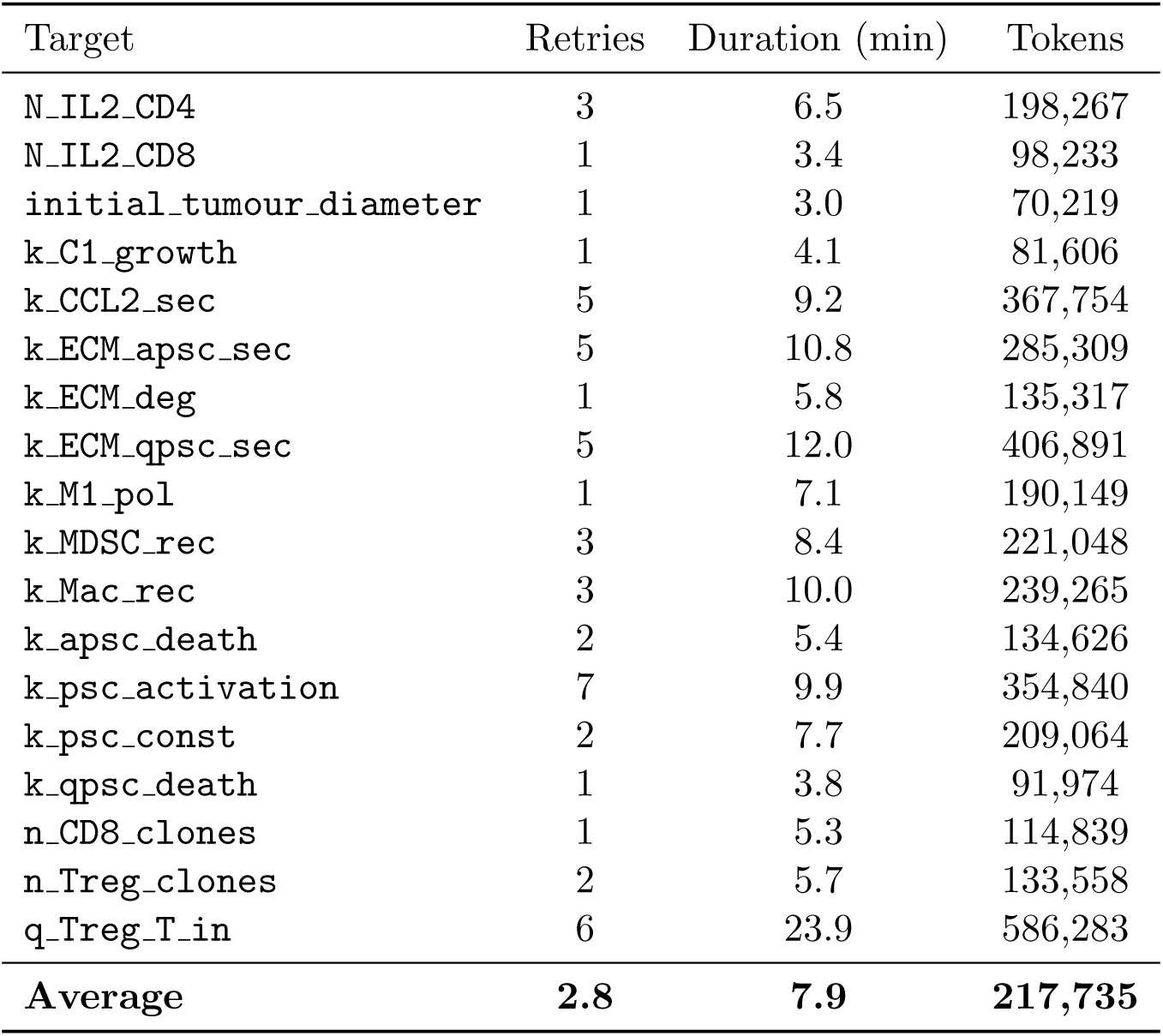
Extraction metrics for the 18 Logfire-instrumented SubmodelTargets. Retries: automated re-extractions triggered by validator failures before any human review. Duration: wall-clock extraction time. Tokens: total LLM tokens consumed. The final row reports the mean across targets.

| Target | Retries | Duration (min) | Tokens |
| --- | --- | --- | --- |
| N_IL2_CD4 | 3 | 6.5 | 198,267 |
| N_IL2_CD8 | 1 | 3.4 | 98,233 |
| initial_tumour_diameter | 1 | 3.0 | 70,219 |
| k_C1_growth | 1 | 4.1 | 81,606 |
| k_CCL2_sec | 5 | 9.2 | 367,754 |
| k_ECM_apsc_sec | 5 | 10.8 | 285,309 |
| k_ECM_deg | 1 | 5.8 | 135,317 |
| k_ECM_qpsc_sec | 5 | 12.0 | 406,891 |
| k_M1_pol | 1 | 7.1 | 190,149 |
| k_MDSC_rec | 3 | 8.4 | 221,048 |
| k_Mac_rec | 3 | 10.0 | 239,265 |
| k_apsc_death | 2 | 5.4 | 134,626 |
| k_psc_activation | 7 | 9.9 | 354,840 |
| k_psc_const | 2 | 7.7 | 209,064 |
| k_qpsc_death | 1 | 3.8 | 91,974 |
| n_CD8_clones | 1 | 5.3 | 114,839 |
| n_Treg_clones | 2 | 5.7 | 133,558 |
| q_Treg_T_in | 6 | 23.9 | 586,283 |
| **Average** | **2.8** | **7.9** | **217,735** |

None of the extractions passed validation on the first attempt; all required at least one automated retry. Seven targets succeeded after a single retry, three each after two, three, and five retries, and one each after six and seven retries; the ECM and CCL2 secretion parameters and PSC activation required as many as 7 iterations to resolve unit ambiguities or conflicting literature. Each extraction took 7.9 minutes on average, consuming 217,735 tokens per target (3.9M total). Across the 50 retries, the validation exceptions captured in tracing (Table 2) were dominated by unit errors, followed by prior specification, then fabrication (invalid DOIs) and code errors. The LLM used 210 web searches and 22 code executions.

**Table 2:** Categories of validation exceptions captured during batch-extraction tracing. The total is a lower bound: it counts distinct exception types per target, so repeated retries for the same kind of error collapse to a single entry, whereas the validators triggered 50 retries in total across the 18 instrumented extractions. Categories match the validator types described in the Validation Framework (Methods); “prior” denotes an uncertainty range incompatible with the stated translation context.

| Error Category | Count | % |
| --- | --- | --- |
| units | 9 | 38% |
| prior | 5 | 21% |
| fabrication | 4 | 17% |
| code | 4 | 17% |
| other | 1 | 4% |
| hallucination | 1 | 4% |
| **Total** | **24** |  |

#### 3.1.1 Source Characteristics

The 37 curated targets drew from diverse sources (Supplementary Section 8): 46% human clinical and 8% human in vitro, with the remainder from animal studies and one review. About 57% used human data directly; the rest required cross-species scaling from mouse (35%) or rat (8%). Fewer than half (46%) came from PDAC-specific experiments; the rest relied on proxy (43%) or related (11%) indications.

### 3.2 Human Curation

To quantify human curation, we compared raw LLM output against final curated versions. All curation was performed interactively with Claude Code (claude-opus-4-6), with the modeler directing edits and modeling decisions while the LLM handled text generation, code writing, and literature comprehension. No batch extraction was used without modification; in a documented review of one batch (12 CalibrationTarget files), the modeler did not promote 58% (7 of 12) to the corpus; recorded reasons included placeholder outputs from LLM tool-access failures, parse errors, observable mislabeling, and source/assumption conflicts. This rate comes from a single small batch and should be read as indicative rather than a precise rejection rate.

Table 3 reports the field-level revision rates for the batch-extracted targets that carry a pre-curation draft. The revisions concentrate in the modeling-judgment fields: forward-model type (65%) and source relevance (every file) for SubmodelTargets, and observable code (39%) and species mapping for CalibrationTargets. Source relevance was revised in fewer batch CalibrationTargets than SubmodelTargets, reflecting their smaller and more curated input pool.

**Table 3:** Modeler revision of batch-extracted targets, by field. Each rate is the fraction of files whose field changed between the raw LLM draft and the curated version, over the cohorts that carry a pre-curation draft: 37 SubmodelTargets and 18 surviving batch-extracted CalibrationTargets (curated from 22 paired drafts, 4 of which were retired during curation). The full CalibrationTarget corpus is these 18 batch-extracted targets plus 27 interactive targets (45 total); the interactive targets were shaped during extraction and have no pre-curation draft to diff against.

| Schema ($n$) | Field | Revised |
| --- | --- | --- |
| SubmodelTarget (37) | forward-model type | 65% |
|  | prior parameters | 46% |
|  | input values | 46% |
|  | source relevance | 100% |
| CalibrationTarget (18) | observable code | 39% |
|  | species list | 28% |
|  | empirical data | 17% |
|  | source relevance | 28% |

A further 27 CalibrationTargets in the final cohort were extracted interactively with Claude Code using source PDFs loaded directly into the session. The modeler provided domain guidance (species mapping, denominator definitions, distribution choices) while the LLM handled literature comprehension and code generation. These files required minimal subsequent revision (a single shared reference value correction across 7,619 total lines); modeler judgment was embedded in the extraction process rather than applied post-hoc.

The curation edits fell into seven recurring categories, including unit conventions, source selection, denominator choices, and scenario-specific structural constraints; the full list is in Supplementary Section 14.1.

The reduced revision rate for interactively extracted CalibrationTargets is observational rather than controlled. The batch pipeline used GPT-5.1 and the interactive pipeline used Claude Opus 4.6; these models differ in context handling and characteristic error profiles, so any per-mode difference may partly reflect model capability rather than collaboration mode. The interactive work also followed curation of the batch outputs, meaning the modeler entered interactive extraction with working knowledge of where the LLM tended to fail on this corpus. The data are consistent with interactive curation reducing post-hoc revision, but a controlled comparison would require the same underlying model in both modes and randomized assignment of targets.

### 3.3 CalibrationTarget Extraction

We applied the CalibrationTarget schema to 45 full-model endpoints for the same PDAC QSP model, organized into three scenario groups: 26 baseline (no treatment) targets capturing the untreated tumor microenvironment, 18 neoadjuvant immunotherapy targets (GVAX ± nivolumab), and 1 clinical progression target (Table 4). All 45 targets pass schema validation including observable code execution, distribution code verification, and unit consistency checks.

**Table 4:** CalibrationTarget summary by scenario group.

| Scenario Group | Targets |
| --- | --- |
| Baseline (no treatment) | 26 |
| Neoadjuvant immunotherapy (GVAX ± nivolumab) | 18 |
| Clinical progression | 1 |
| **Total** | **45** |

Of the 45 targets, 18 were extracted by GPT-5.1 through the batch pipeline and 27 interactively with Claude Opus 4.6 using source PDFs. The 18 batch-extracted figure is the surviving subset of the 22 drafts produced by the batch pipeline; 4 drafts were curated and then retired (Section 3.2). The neoadjuvant immunotherapy targets draw on per-patient data digitized from scatter plots; 15 CalibrationTargets (33% of the total) carry figure-digitized data. Curation metrics for both extraction modes are reported in Section 3.2.

The observables span 25 unique model species, with complexity ranging from 1 to 19 species per target (mean 8.4). The targets drew from 18 unique primary sources (from years 2007–2026). Supplementary Section 9 provides detailed per-target metrics.

Per-target extraction metrics (retry counts, token usage, and validator-error categories) were collected through Logfire instrumentation that was active only for the SubmodelTarget batch run. The CalibrationTarget extraction runs predate that instrumentation, so equivalent per-target metrics are not available for this cohort; the SubmodelTarget metrics in Table 1 are reported as representative of the automated retry loop, and the schema-validation and curation results above are the performance evidence specific to CalibrationTargets.

### 3.4 Code Generation and Inference

All 37 SubmodelTargets were translated to a joint Julia inference script targeting Turing.jl ^25^. The translator identified 19 unique parameters; 11 of these appeared in more than one target, and the translator fit each one against all of its sources at once. Combining several measurements of the same parameter into one prior is normally done by hand; here the shared parameter names let the translator do it automatically, so each posterior draws on every source that measured the parameter, not just one. All parameters converged (*R̂* < 1.01), with bulk-ESS *>* 2600 and tail-ESS *>* 1300 throughout (Table 9). Recognizable quantities fall in clinically measured ranges: the inferred tumor doubling time of ∼130 days is consistent with PDAC volume-doubling-time measurements ^27,28^. Not all estimates are equally constrained: Supplementary Section 10 notes that the IL-2-dependent CD8+ division count falls below its expected range. Full inference results including convergence diagnostics, posterior distributions, and model type breakdown are reported in Supplementary Section 10.

## 4 Discussion

Using MAPLE on the PDAC model, we extracted and curated 37 SubmodelTargets and 45 CalibrationTargets. Every extracted value is tied to a verbatim source snippet and a DOI-resolved citation, every observable carries unit-checked code, and 11 of the 19 unique parameters identified across the corpus appear in more than one target. Per-target manual extraction leaves such constraints scattered across separate documents. MAPLE binds them by shared parameter name, fitting each shared parameter against every source that constrains it in a single inference. The corpus, schemas, validators, prompts, and reference database are pinned at the commit reported in the Data Availability Statement, so the analysis can be re-run from the public repository. This matches the IQ Consortium’s recommendations for explainability and documented provenance in AI-assisted modeling ^7^.

MAPLE demonstrates that structured schemas can serve as a productive collaboration interface between LLMs and modelers for QSP model calibration. The curation analysis shows that LLM-generated drafts are competent at the bibliographic, transcription, and unit-handling tasks the schema encodes, while a substantive minority of fields, concentrated in source selection, prior specification, and forward- or observable-model choice, were revised by the modeler. Rather than automating away the modeler, the framework restructures the collaboration: the schema defines what must be captured, and validators catch errors from both LLM and human sources.

### 4.1 Validation as the Core Mechanism

Across the 18 instrumented SubmodelTarget extractions, none passed validation on the first attempt: the validators triggered 50 retries before any human review, spanning unit errors, fabricated citations, and value-snippet mismatches. The retry loop resolved most issues without human intervention, with extractions requiring multiple retries (up to 7) typically involving ambiguous unit conventions or conflicting literature values. The same automated retry pipeline was used for batch-extracted CalibrationTargets, though per-CT retry counts were not instrumented (the Logfire-instrumented pipeline post-dated the CalibrationTarget extraction runs; the gap is documented in Section 3.3).

The value-in-snippet requirement proved particularly effective, catching a common LLM behavior: paraphrasing source text rather than quoting it exactly. Together, these validators support human review: a reviewer can verify each extraction by checking the snippet against the source paper, without re-reading entire methods sections.

Passing all validators confirms that a target is well-formed, internally consistent, traceable to a resolvable source, dimensionally valid, and executable, but not that the scientific judgment in the calibration layer is correct. Every final SubmodelTarget and CalibrationTarget in this study passes the full validator suite (verified by re-running the Pydantic validator pipeline at MAPLE v0.1.0 against all final YAMLs; the check is scripted as `scripts/verify_validation.sh` in the analysis repository); no record retains an unresolved validator failure. The errors that survive automated validation are therefore judgment errors rather than validator failures: a well-formed forward-model type that is inappropriate for the experiment, a valid prior that is mis-centered, or a resolvable source that is only weakly applicable to the target indication. The curation revision rates in Section 3.2 count these residual errors, which is why curation is necessary even when every final record passes automated validation.

### 4.2 Schema Design Choices

The separation of data extraction from model specification clarifies where human judgment is required (the inputs should be objective; the calibration layer involves modeling choices), enables reuse (if a modeling assumption changes, the inputs remain valid), and supports independent review of data versus modeling decisions. The schema and validators also work without an LLM. A modeler can fill in the fields by hand and still get the provenance trail, the automatic checks, and the route to inference code, and the validators flag transcription, unit, and reference errors no matter who entered the data.

The dual-schema architecture applies the same data-first principles at two scales of calibration. SubmodelTarget operates on isolated experiments that constrain individual parameters through simplified forward models; CalibrationTarget operates on clinical and in vivo endpoints that constrain the integrated model through full-species observables.

### 4.3 Modeler Contribution and Limitations of Extracted Data

The LLM produces usable forward-model choices and observable code when it has the right context, but cannot tell when the context is incomplete or when a paper’s marker definitions disagree with the model’s categorization. The modeler supplies that context and reviews each record for the lapses the validators do not catch. Every claim in the YAML records a source reference and verbatim snippet, and cross-species or cross-indication uses require a documented justification, so the modeler’s input is itself auditable. This is by design: the modeler’s reasoning about which data apply and how uncertain they are is the costly part of calibration, and it usually goes unrecorded. MAPLE writes it down in a form that can be re-run and checked and that outlives any one person on the project. Supplementary Section 14.1 lists the recurring categories of modeler-supplied context observed in this corpus. As LLMs improve, more of the drafting can move to the model, but the schema keeps the modeler’s decisions on the record either way.

The source characteristics reveal a fundamental challenge in QSP calibration: only 46% of SubmodelTargets came from PDAC-specific experiments, and 35% required cross-species translation from mouse data. The schema addresses this through the `source_relevance` block, which requires that translation uncertainty propagate to prior widths.

The token cost reported above (217,735 tokens per SubmodelTarget, averaging roughly $0.37 per target and $6.72 across the 18 instrumented targets at the API pricing in effect during these runs ^29^) is small in absolute terms, so the dominant cost of the workflow is modeler time rather than compute. That time differs by collaboration mode: batch curation requires the modeler to review, and often reject or repair, drafts that may carry unit, source-selection, or denominator errors, while interactive extraction front-loads modeler effort during extraction but leaves little to revise afterwards. We do not report a controlled time comparison against fully manual extraction because the two were not run on the same targets under matched conditions, and net time efficiency depends on the model, the literature base, and the modeler’s familiarity with the framework. MAPLE does not eliminate modeler effort; it records that effort in a form that can be audited and re-run, naming the decisions the modeler made and why, which a free-text extraction does not.

### 4.4 Relationship to Existing Approaches

MAPLE addresses a gap between general-purpose biomedical extraction frameworks^17,18^, which do not address QSP-specific downstream requirements (uncertainty quantification, forward model specification, code generation), and QSP-focused tools like QSP-Copilot^16^, which include literature extraction components but emphasize model structure discovery rather than quantitative parameter calibration with documented provenance and uncertainty propagation. The validation approach combines grounding strategies from retrieval-augmented generation^30^ and constrained decoding ^31^ with domain-specific validators (DOI resolution, unit parsing, code execution) tailored to quantitative scientific extraction.

Unlike knowledge graph approaches that prioritize coverage across thousands of abstracts ^14,32^ and can tolerate some extraction errors, parameter calibration requires higher per-target stringency because each value directly influences model predictions. A more detailed comparison of individual tools and approaches is provided in Supplementary Section 15.

### 4.5 Limitations and Future Work

Several limitations merit discussion. First, the framework relies on the LLM’s built-in web search for source discovery, so extraction quality is bounded by the LLM’s ability to locate and access papers. Closed-access publications may be only partially available (e.g., abstract only), limiting extraction depth. Emerging agentic capabilities (LLM-controlled browsers with institutional authentication) may address this. Work in progress extends the framework to a staged extraction pipeline that retrieves full-text papers in advance and batches extraction over them systematically, which would make the external snippet-in-paper check part of the automated gate rather than an on-demand step. The framework also depends on evolving LLM capabilities: extraction is stochastic, and performance characteristics vary across providers and model versions.

Second, the framework still needs substantial modeler input. For SubmodelTargets, the modeler changed the forward model type in 65% of files and adjusted prior parameters in 46%. Prompt improvements could reduce some recurring errors, but the underlying scientific decisions are likely to remain with the modeler even as LLM capabilities improve. CalibrationTargets required different curation: observable code (changed in 39% of batch-extracted files), Monte Carlo distribution derivation, and denominator audit fields demand modeling expertise that current LLMs provide as a starting point.

Third, extracting structured quantitative data from figures remains challenging; the neoadjuvant immunotherapy targets required manual digitization of scatter plots. Integrating multimodal LLM vision capabilities is a natural direction for future work. Fourth, per-target extraction metrics are available only for the SubmodelTarget cohort, as discussed in Section 3.3.

#### Generalization requirements

This evaluation covers one QSP model in one disease area, extracted and curated by one modeling group, so it characterizes the framework rather than measuring how broadly it generalizes. Several components are domain-independent: the separation of an inputs layer from a calibration layer, the typed-role fields, the value-in-snippet, DOI-resolution, and code-execution validators, and the provenance requirements all apply to any literature-derived parameter. The forward-model type library is the main domain-specific surface. 49% of SubmodelTargets used the generic algebraic type, which indicates that the 15 built-in templates covered roughly half of the use cases even within this model; a portion of those algebraic uses are genuinely simple derived relations (half-lives, ratios) for which no structured template is needed, but the fraction also shows the template library is not exhaustive. A non-oncology application, for instance a PK/PD model, would likely require additional templates for compartmental pharmacokinetics, receptor occupancy, or indirect-response dynamics; the generic algebraic, direct-fit, and custom fallbacks accept such cases today but without the structured role typing that the built-in templates provide. The model-aware search and shared-parameter-name binding assume a structured QSP model is already in hand, which holds in any domain but means the framework supports calibration of an existing model rather than model construction.

## Study Highlights

### What is the current knowledge on the topic?

QSP model calibration requires literature data spanning experimental and clinical endpoints. Manual curation is labor-intensive and inconsistently documented, while LLM extraction hallucinates values and fabricates citations.

### What question did this study address?

Can structured schemas serve as a collaboration interface between LLMs and modelers for QSP calibration, and what is each party’s contribution?

### What does this study add to our knowledge?

Using MAPLE on a PDAC QSP model, we curated 37 SubmodelTargets and 45 CalibrationTargets, every value tied to a verbatim snippet and a resolved DOI; 11 of 19 parameters appear in more than one target. Automated validators rejected every extraction at least once (50 retries across 18 instrumented SubmodelTargets) before human review, and modeler curation supplied the remaining scientific context.

### How might this change drug discovery, development, and/or therapeutics?

The framework produces reproducible calibration data: every value traceable to its source and uncertainty, priors built from several sources, and the modeler’s reasoning recorded so it is not lost when personnel change. A reviewer can then check the calibration data directly rather than take it on trust.

## Acknowledgments

Claude Opus 4.6 (Anthropic, model identifier `claude-opus-4-6`) and GPT-5.1 (OpenAI, model identifier `gpt-5.1`) were used for literature search, data extraction, and code generation as described in the Methods, and Claude Code (`claude-opus-4-6`) assisted with manuscript preparation and interactive curation. All AI-generated content was reviewed and verified by the authors.

## Author Contributions

J.E. designed research, performed research, analyzed data, and wrote the manuscript. A.S.P. designed research and wrote the manuscript.

## Data Availability Statement

MAPLE (the SubmodelTarget and CalibrationTarget schemas and validation framework, plus the inference code generator in the v0.1.0 release) is available at https://github.com/popellab/maple. The results reported here were generated with MAPLE v0.1.0 (commit `7f1faa4`), and the repository includes the representative extraction prompts used for both schemas. Extraction inputs, curated YAML targets, and scripts to reproduce all paper statistics are available at https://github.com/popellab/maple-paper. The 37 SubmodelTargets and 45 CalibrationTargets used in this study are provided as supplementary YAML files. Generated Julia inference scripts are included in the supplementary materials. The underlying PDAC QSP model is part of separate work that has not yet been published; the structural representation that MAPLE consumes (model structure, species and parameter definitions, and unit mappings) is included in the `maple-paper` repository, so the extraction inputs and reported evaluation are fully reproducible. The complete PDAC model will be released with its own publication.

