## Supplement for "Structured Schemas for Provenance-Rich, LLM-Assisted QSP Model Calibration"

**11 Model-Aware Prompt Construction Details** **33**

**12 Schema Implementation Details** **36**

**13 Source Relevance Assessment Details** **36**

**14 Collaboration Mode Details** **37**
14.1 Categories of modeler-supplied context . . . 38

**15 Detailed Comparison to Existing Approaches** **38**

### 1 Complete SubmodelTarget Example

This section presents a complete SubmodelTarget for activated pancreatic stellate cell (aPSC) proliferation, demonstrating all schema components.

#### 1.1 Overview

This target constrains the proliferation rate parameter `k_apsc_prolif` using data from Schneider et al. (2001), who measured PSC proliferation in response to PDGF stimulation via BrdU incorporation assay.

#### 1.2 Complete YAML Specification

```yaml
# Schema: SubmodelTarget
# Target: Activated PSC proliferation rate

target_id: psc_proliferation_PDAC_deriv001

# =============================================================================
# INPUTS: Extracted values with full provenance
# =============================================================================
inputs:
  - name: pdgf_fold_mean
    value: 4.37
    units: dimensionless
    input_type: direct_measurement
    role: target
    source_ref: Schneider2001_PSC_PDGF
    source_location: Abstract
    value_snippet: "Cell proliferation (4.37 +/- 0.49- and 2.96 +/-
                    0.39-fold of control)."

  - name: pdgf_fold_sd
    value: 0.49
    units: dimensionless
    input_type: direct_measurement
    role: auxiliary
    source_ref: Schneider2001_PSC_PDGF
    source_location: Abstract
    value_snippet: "Cell proliferation (4.37 +/- 0.49- and 2.96 +/-
                    0.39-fold of control)."

# =============================================================================
# CALIBRATION: Everything needed for inference
# =============================================================================
calibration:
  parameters:
    - name: k_apsc_prolif
```

```
    units: 1/day
    prior:
      distribution: lognormal
      mu: 0.0      # log(1.0) - expect ~1/day for 4x fold-change in 1 day
      sigma: 1.0
      rationale: |
        Wide log-normal prior centered at 1/day. A 4-fold increase in
        1 day implies k = ln(4) = 1.4/day, so prior median of 1/day
        is reasonable.

forward_model:
  type: exponential_growth
  rate_constant: k_apsc_prolif
  data_rationale: |
    BrdU incorporation assay measures proliferation as fold-change
    over ~24 hours under constant PDGF stimulus. With a single high
    PDGF dose and short duration, exponential growth (dN/dt = k*N)
    fits the data structure.
  submodel_rationale: |
    The full QSPIO_PDAC model has aPSC proliferation via:
    k_apsc_prolif * aPSC * (PDGF / (PDGF_50_prolif + PDGF)).
    At 50 ng/mL PDGF >> PDGF_50_prolif, the saturation term ~ 1,
    reducing to exponential growth with rate k_apsc_prolif.
  independent_variable:
    name: time
    units: day
    span: [0.0, 1.0]
    rationale: |
      Proliferation assay duration assumed to be ~24 hours based on
      typical cytokine incubation protocols for rat PSCs.
  state_variables:
    - name: aPSC_rel
      units: dimensionless
      initial_condition:
        value: 1.0
        rationale: |
          PSC proliferation data are reported as fold of control. We
          normalize the initial aPSC population to 1.0 (100% control)
          to match this convention.

error_model:
  - name: proliferation_fold_change
    observable:
      type: identity
      state_variables: [aPSC_rel]
      rationale: |
        The state variable aPSC_rel is the fold-change in aPSC number
        relative to baseline (control = 1.0), which directly matches
```

```
     the reported proliferation fold-change.
   units: dimensionless
   uses_inputs:
    - pdgf_fold_mean
    - pdgf_fold_sd
   evaluation_points: [1.0]
   sample_size: 3
   sample_size_rationale: |
    Sample size not explicitly reported; assumed n=3 independent
    experiments, typical for in vitro PSC BrdU proliferation assays.
   distribution_code: |
    def derive_distribution(inputs, ureg):
      import numpy as np

      rng = np.random.default_rng(42)

      mean = inputs['pdgf_fold_mean'].magnitude
      sd = inputs['pdgf_fold_sd'].magnitude
      n = 10000

      # Use lognormal for positive fold-changes
      mu_log = np.log(mean**2 / np.sqrt(mean**2 + sd**2))
      sigma_log = np.sqrt(np.log(1 + sd**2 / mean**2))

      samples = rng.lognormal(mu_log, sigma_log, n)

      median_val = np.median(samples)
      ci_lower = np.percentile(samples, 2.5)
      ci_upper = np.percentile(samples, 97.5)

      return {
        'median': [median_val],
        'ci95': [[ci_lower, ci_upper]],
        'units': 'dimensionless'
      }
   likelihood:
    distribution: lognormal
    rationale: |
     Fold-changes are positive and multiplicative, making
     lognormal appropriate.

 identifiability_notes: |
  With a single timepoint (~24h) and normalized initial condition
  (aPSC_rel(0)=1), only k_apsc_prolif is estimated. If substantial
  PSC death occurred over the assay interval, the inferred rate
  would represent net growth rather than pure proliferation.

# ========================================================================
```

```yaml
# EXPERIMENTAL CONTEXT
# ============================================================================
experimental_context:
  species: rat
  system: in_vitro_primary_cells
  indication: PDAC
  cell_types:
    - name: pancreatic stellate cell
      phenotype: activated (alpha-SMA positive)
      isolation_method: Isolated from normal Wistar rat pancreas,
                        activated by culture on plastic
  culture_conditions:
    culture_type: 2d_monolayer

# ============================================================================
# STUDY INTERPRETATION
# ============================================================================
study_interpretation: |
  Schneider et al. (2001) measured proliferation of cultured rat
  pancreatic stellate cells (PSCs) in response to growth factors using
  BrdU incorporation. Addition of 50 ng/mL PDGF increased PSC
  proliferation to 4.37 +/- 0.49-fold of control, indicating that PDGF
  acts as a strong mitogen for activated PSCs. We interpret the
  50 ng/mL PDGF condition as approximating a near-maximal PDGF stimulus
  and use the reported fold-change over ~24 hours as a quantitative
  constraint on the maximal PDGF-driven proliferation rate parameter
  k_apsc_prolif.

key_assumptions:
  - |
    50 ng/mL PDGF corresponds to a near-saturating concentration for
    PSC proliferation, so PDGF/(PDGF_50_prolif + PDGF) ~ 1.
  - |
    The proliferation assay duration is approximately 24 hours,
    consistent with typical cytokine incubation protocols.
  - |
    BrdU incorporation fold-change is proportional to fold-change in
    PSC cell number (negligible death over 24 hours).
  - |
    Rat activated PSC proliferation kinetics in vitro provide an upper
    bound for human tumor aPSC proliferation in vivo.

key_study_limitations:
  - |
    Proliferation was measured via BrdU incorporation rather than
    direct cell counts.
  - |
    Single PDGF concentration and single assay interval constrain
```

```
    only the maximal rate.
  - |
    Data from rat PSCs in vitro, not human PDAC-associated stellate cells.
  - |
    Background serum proliferative signals not explicitly modeled.

# ============================================================================
# DATA SOURCES
# ============================================================================
primary_data_source:
  doi: "10.1152/ajpcell.2001.281.2.C532"
  title: "Identification of mediators stimulating proliferation and
          matrix synthesis of rat pancreatic stellate cells"
  authors: [Schneider, Schmid, Liptay, Schmid, Pohl]
  year: 2001
  source_tag: Schneider2001_PSC_PDGF

secondary_data_sources:
  - doi: "10.1136/gut.44.4.534"
    title: "Pancreatic stellate cells are activated by proinflammatory
            cytokines: implications for pancreatic fibrogenesis"
    authors: [Apte, Haber, Darby, Rodgers, McCaughan, Korsten, Pirola, Wilson]
    year: 1999
    source_tag: Apte1999_PSC_Cytokines
    contribution: |
      Provides context on PSC activation and cytokine response protocols.
```

##### 1.3 Schema Structure Summary

The SubmodelTarget schema has the following top-level structure:

- **target_id**: Unique identifier for the target
- **inputs**: List of values extracted from literature with provenance
- **calibration**: Everything needed for inference code generation
  - `parameters`: Parameters to estimate with priors
  - `forward_model`: Forward model specification (built-in type or custom). For ODE models, contains nested `state_variables` (state variables with initial conditions) and `independent_variabl` (e.g. time)
  - `error_model`: Observable specifications mapping model predictions to data with likelihoods
  - `identifiability_notes`: Discussion of parameter identifiability
- **experimental_context**: Species, system, cell types, culture conditions
- **study_interpretation**: Scientific interpretation of the data
- **key_assumptions**: Assumptions made in using this data

- **key_study_limitations**: Known limitations of the source study
- **primary_data_source**: Primary literature reference with DOI
- **secondary_data_sources**: Additional references

### 2 Supported Model Types

The SubmodelTarget schema supports 15 built-in forward model types, organized into four categories.

Table 1: Built-in forward model types organized by category.

| Category | Model Type | Formulation | Key Fields |
|---|---|---|---|
| ODE templates | `exponential_growth` | $dy/dt = k \cdot y$ | `rate_constant` |
| | `first_order_decay` | $dy/dt = -k \cdot y$ | `rate_constant` |
| | `logistic` | $dy/dt = k \cdot y(1 - y/K)$ | `rate_constant`, `carrying_capacity` |
| | `saturation` | $dy/dt = k(1 - y)$ | `rate_constant` |
| | `two_state` | $dA/dt = -kA$, $dB/dt = kA$ | `forward_rate` |
| | `michaelis_menten` | $dy/dt = -V_{\max} y/(K_m + y)$ | `vmax`, `km` |
| Structured steady-state | `steady_state_density` | rate = density × loss | `target_rate`, `observed_density`, `loss_rate` |
| | `steady_state_fraction` | rate $= f \times \rho_{\text{parent}} \times$ loss | `target_rate`, `observed_fraction`, `parent_density` |
| | `steady_state_concentration` | sec $= C \times k_{\text{cl}} \times V$ | `target_rate`, `observed_concentration`, `clearance_rate` |
| | `steady_state_ratio` | $r = k_1/k_2$ | `target_rate`, `observed_ratio`, `opposing_rate` |
| | `steady_state_proliferation` | $f = k_p/(k_p + k_d)$ | `target_rate`, `observed_fraction`, `death_rate` |
| Accumulation | `batch_accumulation` | $C = (\text{sec} \times n \times t)/V$ | `target_rate`, `cell_count`, `incubation_time` |
| Generic/ fallback | `algebraic` | (closed-form formula) | `code`, `code_julia` |
| | `direct_fit` | (curve fitting) | `curve` |
| | `custom` | (user-provided ODE) | `code`, `code_julia` |

Each structured steady-state and accumulation model declares its fields as typed roles: a field can reference a calibration parameter (to be estimated), an extracted input (from the inputs layer), a curated reference value from a shared database, or a numeric literal. This role-based design allows the same template to serve different estimation scenarios depending on which field is treated as the unknown.

### 3 Validation Checks

The validation framework includes several dozen Pydantic `@model_validator` and `@field_validator` methods across the SubmodelTarget and CalibrationTarget schemas. They run after field parsing and raise a `ValidationError` when constraints are violated. The individual validators are fine-grained, but their behaviour is grouped into the ten categories below.

#### 3.1 Summary of Validators

1. **Input reference validation**: All `uses_inputs` references match defined input names
2. **Source reference validation**: All `source_ref` values match defined source tags
3. **Parameter reference validation**: Model parameter roles reference defined parameters
4. **State variable reference validation**: Observable `state_variables` match defined state variables
5. **DOI resolution**: All DOIs resolve via CrossRef API
6. **Value-in-snippet validation**: Extracted values appear verbatim in quoted snippets
7. **Unit validation**: All unit strings parse with Pint
8. **Code syntax validation**: Custom code passes Python AST parsing
9. **Span ordering validation**: Time spans have start < end
10. **ODE model requirements**: ODE models have required state variables and spans

#### 3.2 DOI Resolution Validator

The DOI validator queries the CrossRef API to verify that every cited DOI resolves to a valid publication. This catches fabricated citations before they enter the calibration pipeline. The validator also performs fuzzy matching on title and year to detect metadata inconsistencies.

```
def resolve_doi(doi: str) -> Optional[dict]:
    """Resolve DOI and get metadata from CrossRef."""
    doi_clean = doi.replace("https://doi.org/", "").strip()

    url = f"https://doi.org/{doi_clean}"
    headers = {"Accept": "application/vnd.citationstyles.csl+json"}
    response = requests.get(url, headers=headers, timeout=10)

    if response.status_code != 200:
```

```
        return None  # DOI does not resolve

    metadata = response.json()
    return {
        "title": metadata.get("title", ""),
        "first_author": metadata.get("author", [{}])[0].get("family", ""),
        "year": metadata.get("issued", {}).get("date-parts", [[]])[0][0]
    }

@model_validator(mode="after")
def validate_doi_resolution_and_metadata(self) -> "SubmodelTarget":
    """Validate that DOIs resolve and metadata matches."""
    errors = []

    metadata = resolve_doi(self.primary_data_source.doi)
    if metadata is None:
        errors.append(f"DOI '{self.primary_data_source.doi}' failed to resolve")
    else:
        # Check title match (fuzzy)
        if not fuzzy_match(self.primary_data_source.title, metadata["title"]):
            errors.append(f"Title mismatch with CrossRef")
        # Check year match
        if self.primary_data_source.year != metadata["year"]:
            errors.append(f"Year mismatch: {self.primary_data_source.year} vs {metadata['year']

    if errors:
        raise ValueError("DOI validation errors:\n  - " + "\n  - ".join(errors))
    return self
```

**Example failure**: If an LLM fabricates a citation with DOI `10.1234/fake.2023.12345`, the CrossRef query returns a 404 error and validation fails with:

```
ValidationError: DOI '10.1234/fake.2023.12345' failed to resolve.
Verify at https://doi.org/10.1234/fake.2023.12345
```

#### 3.3 Value-in-Snippet Validator

The value-in-snippet validator checks that each extracted numeric value appears in its associated source text. This catches hallucinations where an LLM extracts a plausible but incorrect value. The validator handles multiple numeric formats including scientific notation, percentages, and Unicode superscripts.

```
def check_value_in_text(text: str, value: float) -> bool:
    """Check if numeric value appears in text."""
    text_norm = text.lower().replace(",", "")
    patterns = []

    # Direct value and common formats
    patterns.append(str(value))
```

```
    patterns.append(f"{value:.1f}")
    patterns.append(f"{value:.2f}")

    # Scientific notation (e.g., "1.5e-3", "1.5E-03")
    if abs(value) < 0.01 or abs(value) > 1000:
        patterns.append(f"{value:e}")
        patterns.append(f"{value:.2e}")

    # Percentage format (if value is between 0 and 1)
    if 0 < value < 1:
        pct = value * 100
        patterns.extend([f"{pct}%", f"{pct:.0f}%", f"{pct:.1f}%"])

    return any(p.lower() in text_norm for p in patterns)

@model_validator(mode="after")
def validate_input_values_in_snippets(self) -> "SubmodelTarget":
    """Validate that extracted values appear in their value_snippet."""
    errors = []
    for inp in self.inputs:
        if not inp.value_snippet:
            continue
        # Skip types that may not have explicit values in text
        if inp.input_type in (InputType.INFERRED_ESTIMATE, InputType.ASSUMED_VALUE):
            continue

        if not check_value_in_text(inp.value_snippet, inp.value):
            errors.append(
                f"Input '{inp.name}': value {inp.value} not found in snippet "
                f"'{inp.value_snippet[:60]}...'"
            )

    if errors:
        raise ValueError("Value-snippet mismatches (possible hallucination):\n  - "
                         + "\n  - ".join(errors))
    return self
```

**Example failure**: If an input claims `value: 4.37` but the snippet reads "Cell proliferation was 3.21-fold of control", validation fails with:

```
ValidationError: Value-snippet mismatches (possible hallucination):
  - Input 'pdgf_fold_mean': value 4.37 not found in snippet
    'Cell proliferation was 3.21-fold of control...'
```

This validator is particularly effective because LLMs that hallucinate values rarely fabricate consistent snippets, so the mismatch provides a reliable detection signal.

#### 3.4 Unit Validation

The unit validator parses all unit strings using Pint, a Python library for physical quantities. Malformed or unrecognized units trigger validation failure.

```
@model_validator(mode="after")
def validate_units_are_valid_pint(self) -> "SubmodelTarget":
    """Validate that all unit strings are valid Pint units."""
    from pint import UnitRegistry
    ureg = UnitRegistry()
    errors = []

    def check_unit(unit_str: str, location: str):
        try:
            ureg(unit_str)
        except Exception as e:
            errors.append(f"{location}: ’{unit_str}’ is not valid ({e})")

    for inp in self.inputs:
        check_unit(inp.units, f"Input ’{inp.name}’")
    for param in self.calibration.parameters:
        check_unit(param.units, f"Parameter ’{param.name}’")

    if errors:
        raise ValueError("Invalid Pint units:\n  - " + "\n  - ".join(errors))
    return self
```

**Example failure**: If a parameter specifies `units: cells/mL/day` (invalid compound unit), validation fails with:

```
ValidationError: Invalid Pint units:
  - Parameter ’k_recruit’: ’cells/mL/day’ is not valid (...)
```

#### 3.5 Reference Consistency Validators

Four validators check that all cross-references within the schema are consistent:

- **Input references**: Every `uses_inputs` entry and `initial_condition.input_ref` must match a defined input name.
- **Source references**: Every `source_ref` must match `primary_data_source.source_tag` or a `secondary_data_sources[].source_tag`.
- **Parameter references**: When a model field like `rate_constant` contains a string (not an `InputRef`), it must match a parameter name in `calibration.parameters`.
- **State variable references**: Every state variable referenced in `observable.state_variables` must exist in `calibration.state_variables`.

These validators catch internal inconsistencies where an LLM generates references to entities that don’t exist elsewhere in the document.

##### 3.6 Code Execution Validation

Code validation goes beyond syntax checking. The validator parses the code structure, then executes it with mock data to verify correct behavior. For `observation_code`, execution must return the required dictionary keys (`value`, `sd`, `sd_uncertain`) with appropriate types. For observable code, execution must return arrays with correct units and dimensions. This catches logic errors and signature mismatches before inference.

##### 3.7 Hardcoded Constant Detection

Hardcoded constant detection scans code blocks for numeric literals that should be documented inputs. The parser identifies constants (e.g., `1440` for a minutes-to-days conversion) that are not in the allowed set (small integers 0–5, common fractions, statistical percentiles). Each flagged constant must either be moved to an input with provenance or documented as an `observable_constant` with a `biological_basis` explanation.

##### 3.8 Scale Consistency Validation

Scale consistency validation executes observable code with mock species data spanning a 1000-fold range. If the output is constant despite varying inputs, or if the output magnitude suggests a ratio/score mismatch (e.g., values near 100 when 0–1 is expected), validation warns of potential unit or scaling errors.

##### 3.9 Control Character Validation

Control character validation scans all string fields for characters that corrupt YAML parsing (ASCII control codes, non-breaking spaces, and similar). PDF copy-paste often introduces invisible characters that cause parsing failures; this validator catches them before they propagate.

##### 3.10 CalibrationTarget-Specific Validation

CalibrationTarget adds validators for full-model observables. Observable code is executed against mock species data to verify correct function signatures, output dimensionality, and unit consistency. A denominator audit requires that density or fraction observables explicitly document what tissue area or cell population serves as the denominator, both experimentally and in the model, catching a common source of order-of-magnitude errors. Species existence checks verify that every species referenced in the observable code exists in the model structure. Distribution code is executed with the declared inputs to verify that computed summary statistics (median and CI95) match the values reported in the empirical data block within tolerance (1% for median, 10% for CI bounds), catching transcription errors and code bugs before they propagate to calibration.

#### 4 Generated Julia Code Structure

The translator generates Julia scripts with the following structure:

```
using DifferentialEquations
using Turing
using Distributions

```

```
# Observed data (from inputs layer)
const obs_proliferation_median = 4.37
const obs_proliferation_sigma = 0.25  # derived from CI95

# ODE function
function ode_proliferation!(du, u, p, t)
    k = p[1]
    du[1] = k * u[1]  # exponential growth
end

# Simulate function
function simulate_proliferation(k; t_span=(0.0, 1.0), u0=[1.0])
    prob = ODEProblem(ode_proliferation!, u0, t_span, [k])
    sol = solve(prob, Tsit5())
    return sol[end][1]  # final value
end

# Turing model
@model function proliferation_model()
    # Prior
    k_apsc_prolif ~ LogNormal(0.0, 1.0)

    # Simulate
    pred = simulate_proliferation(k_apsc_prolif)

    # Likelihood
    obs_proliferation_median ~ LogNormal(log(pred), obs_proliferation_sigma)
end

# Run inference
model = proliferation_model()
chain = sample(model, NUTS(), MCMCThreads(), 1000, 4)
```

For joint inference over multiple targets, the translator identifies shared parameters by name and generates a single model with one prior per unique parameter.

### 5 Pydantic Model Definitions

This section provides the key Pydantic model definitions that implement both schemas. The full implementations are available in the repository at `src/qsp_llm_workflows/core/calibration/`.

#### 5.1 Top-Level Model

The `SubmodelTarget` class is the entry point for validation. When a YAML file is loaded, Pydantic recursively validates all nested models.

```
class SubmodelTarget(BaseModel):
    """
```

```
    Submodel-based calibration target for QSP parameter inference.

    Separates:
    - ‘inputs‘: What was extracted from papers (with full provenance)
    - ‘calibration‘: How to use those inputs for inference
    """
    target_id: str = Field(description="Unique identifier for this target")

    # Extracted data with provenance
    inputs: List[Input] = Field(
        description="Values extracted from literature with full provenance"
    )

    # Calibration specification
    calibration: Calibration = Field(
        description="Everything needed for inference code generation"
    )

    # Experimental context
    experimental_context: ExperimentalContext = Field(
        description="Experimental context for the target"
    )

    # Study interpretation
    study_interpretation: str = Field(
        description="Scientific interpretation of how the study data informs the model"
    )
    key_assumptions: List[str] = Field(
        description="Key assumptions made in using this data"
    )
    key_study_limitations: Optional[List[str]] = Field(
        default=None, description="Known limitations of the study"
    )

    # Data sources
    primary_data_source: PrimaryDataSource = Field(
        description="Primary literature source"
    )
    secondary_data_sources: Optional[List[SecondaryDataSource]] = Field(
        default=None, description="Additional literature sources"
    )

    # Validators (10 total) defined via Pydantic decorators
    # See Section S3 for implementation details
```

#### 5.2 Input Model

Each input captures a single value extracted from literature with full provenance:

```python
class InputType(str, Enum):
    """Type of input extracted from literature."""
    DIRECT_MEASUREMENT = "direct_measurement"   # Value reported directly
    PROXY_MEASUREMENT = "proxy_measurement"     # Requires conversion
    EXPERIMENTAL_CONDITION = "experimental_condition"  # Protocol choice
    INFERRED_ESTIMATE = "inferred_estimate"     # Interpreted from qualitative text
    ASSUMED_VALUE = "assumed_value"             # Domain knowledge, not in source

class InputRole(str, Enum):
    """Role of input in calibration."""
    INITIAL_CONDITION = "initial_condition"  # IC for ODE integration
    TARGET = "target"                        # Calibration target (likelihood term)
    FIXED_PARAMETER = "fixed_parameter"      # Fixed value in model
    AUXILIARY = "auxiliary"                  # Supporting data

class Input(BaseModel):
    """A value extracted from literature with full provenance."""
    name: str = Field(description="Unique identifier for this input")
    value: float = Field(description="Extracted numeric value")
    units: str = Field(description="Units of the value")
    uncertainty: Optional[Uncertainty] = Field(default=None)
    n: Optional[int] = Field(default=None, description="Sample size")
    input_type: InputType
    role: Optional[InputRole] = Field(default=None)
    source_ref: str = Field(description="Reference to source_tag in data sources")
    source_location: str = Field(description="Location within the source")
    value_snippet: Optional[str] = Field(
        default=None,
        description="Exact text from paper containing the value (for validation)"
    )
```

##### 5.3 Model Type Discriminated Union

The schema uses Pydantic's discriminated union pattern to support multiple model types, each with type-specific required fields:

```python
class BaseForwardModelSpec(BaseModel):
    """Base class for all forward model specifications."""
    data_rationale: str = Field(
        description="Why this model type fits the experimental data"
    )
    submodel_rationale: str = Field(
        description="Why this is a valid submodel of the full QSP model"
    )

class ExponentialGrowthModel(BaseForwardModelSpec):
    """Exponential growth: dy/dt = k * y"""
    type: Literal["exponential_growth"] = "exponential_growth"
```

```
    rate_constant: ParameterRole = Field(
        description="Rate constant parameter name or input_ref"
    )

class TwoStateModel(BaseForwardModelSpec):
    """Two-state transition: A -> B with first-order kinetics."""
    type: Literal["two_state"] = "two_state"
    forward_rate: ParameterRole = Field(
        description="Forward transition rate constant"
    )

class CustomODEModel(BaseForwardModelSpec):
    """Custom ODE with user-provided code."""
    type: Literal["custom_ode"] = "custom_ode"
    code: str = Field(description="Python ODE function")
    code_julia: str = Field(description="Julia ODE function for inference")

# Discriminated union selects model class based on 'type' field
ForwardModel = Annotated[
    Union[
        FirstOrderDecayModel,
        ExponentialGrowthModel,
        LogisticModel,
        MichaelisMentenModel,
        TwoStateModel,
        SaturationModel,
        SteadyStateDensityModel,
        SteadyStateFractionModel,
        SteadyStateConcentrationModel,
        SteadyStateRatioModel,
        SteadyStateProliferationIndexModel,
        BatchAccumulationModel,
        AlgebraicModel,
        DirectFitModel,
        CustomODEModel,
    ],
    Field(discriminator="type"),
]
```

The `discriminator="type"` annotation tells Pydantic to inspect the `type` field in the YAML to determine which model class to instantiate. This ensures that an `exponential_growth` model is validated against `ExponentialGrowthModel` (which requires `rate_constant`) while a `two_state` model is validated against `TwoStateModel` (which requires `forward_rate`).

##### 5.4 Prior Specification

Priors are specified with distribution type and parameters:

```
class PriorDistribution(str, Enum):
```

```
    """Supported prior distribution types."""
    LOGNORMAL = "lognormal"    # For positive parameters (rates, densities)
    NORMAL = "normal"          # For unconstrained parameters
    UNIFORM = "uniform"        # For bounded parameters
    HALF_NORMAL = "half_normal"  # For positive parameters with mode at 0

class Prior(BaseModel):
    """Prior distribution specification for Bayesian inference."""
    distribution: PriorDistribution
    mu: Optional[float] = Field(default=None, description="Location parameter")
    sigma: Optional[float] = Field(default=None, description="Scale parameter")
    lower: Optional[float] = Field(default=None, description="Lower bound")
    upper: Optional[float] = Field(default=None, description="Upper bound")
    rationale: Optional[str] = Field(default=None)

    @model_validator(mode="after")
    def validate_prior_params(self) -> "Prior":
        """Validate that required parameters are provided for each distribution."""
        if self.distribution == PriorDistribution.LOGNORMAL:
            if self.mu is None or self.sigma is None:
                raise ValueError("lognormal prior requires mu and sigma")
        elif self.distribution == PriorDistribution.UNIFORM:
            if self.lower is None or self.upper is None:
                raise ValueError("uniform prior requires lower and upper")
            if self.lower >= self.upper:
                raise ValueError("uniform lower must be < upper")
        return self
```

##### 5.5 Loading and Validating YAML Files

To load and validate a target:

```
import yaml
from maple.core.calibration import SubmodelTarget

# Load YAML file
with open("psc_proliferation_PDAC_deriv001.yaml") as f:
    data = yaml.safe_load(f)

# Parse and validate (raises ValidationError on failure)
target = SubmodelTarget(**data)

# Access validated fields
print(f"Target: {target.target_id}")
print(f"Parameters: {[p.name for p in target.calibration.parameters]}")
print(f"Model type: {target.calibration.model.type}")
```

If validation fails, Pydantic raises a `ValidationError` with detailed error messages:

```
pydantic_core._pydantic_core.ValidationError: 2 validation errors for SubmodelTarget
calibration.measurements.0.uses_inputs.0
  Input 'typo_input_name' not found in inputs [...]
primary_data_source.doi
  DOI '10.1234/fake' failed to resolve [...]
```

##### 5.6 CalibrationTarget Model

The `CalibrationTarget` schema represents clinical and in vivo endpoints for full-model calibration. Where SubmodelTarget isolates a single mechanism, CalibrationTarget operates on the full model state.

```python
class CalibrationTarget(BaseModel):
    """
    Calibration target for full-model Bayesian inference.

    Represents a clinical or in vivo measurement that constrains
    model behavior at the system level.
    """
    # Observable: how to compute the measurement from model species
    observable: Observable = Field(
        description="Maps full model species to measured quantity"
    )

    # Empirical data: literature-derived distribution
    empirical_data: CalibrationTargetEstimates = Field(
        description="Observed values derived from literature via Monte Carlo"
    )

    # Clinical/experimental context
    experimental_context: ExperimentalContext = Field(
        description="Species, system, indication, treatment, staging"
    )
    scenario: Optional[Scenario] = Field(
        default=None,
        description="Treatment scenario for intervention studies"
    )

    # Interpretation and provenance (shared with SubmodelTarget)
    study_interpretation: str
    key_assumptions: List[str]
    key_study_limitations: Optional[List[str]] = None
    primary_data_source: PrimaryDataSource
    secondary_data_sources: Optional[List[SecondaryDataSource]] = None
    source_relevance: Optional[SourceRelevanceAssessment] = None

    # 30+ CalibrationTarget validators
    # (categorized in Section S3 alongside SubmodelTarget validators)
```

##### 5.7 Observable

The `Observable` specifies how to compute the experimental measurement from full model species. Unlike SubmodelTarget's forward model (which defines isolated dynamics), the observable operates on the solved full-model state.

```
class Observable(BaseModel):
    """
    Full-model observable for CalibrationTarget.

    The code field contains a Python function that computes the
    measured quantity from model species at each time point.
    """
    code: str = Field(
        description="Python: compute_observable(time, species_dict, "
                    "constants, ureg) -> Quantity array"
    )
    units: str = Field(description="Units of the observable output")
    species: List[str] = Field(
        description="Model species accessed (e.g., ['V_T.CD8', 'V_T.C1'])"
    )
    constants: Optional[List[ObservableConstant]] = Field(
        default=None,
        description="Named constants with provenance"
    )
    inputs: Optional[List[SubmodelInput]] = Field(
        default=None,
        description="Literature values used in the observable"
    )
    support: SupportType = Field(
        description="Mathematical domain: positive, non_negative, "
                    "unit_interval, real"
    )
    mapping_rationale: str = Field(
        description="How the literature measurement maps to model species"
    )
    # Denominator audit fields for density/fraction observables
    experimental_denominator: Optional[str] = None
    model_denominator_species: Optional[List[str]] = None
```

Each `ObservableConstant` requires a `biological_basis` explanation and must trace to either the curated reference database or a specific literature source.

##### 5.8 CalibrationTargetEstimates

The empirical data block derives summary statistics from literature inputs via Monte Carlo simulation.

```
class CalibrationTargetEstimates(BaseModel):
```

```
    """
    Empirical data block: literature values -> distribution for likelihood.
    """
    median: List[float] = Field(
        description="Point estimates (length 1 for scalar, "
                    "or matching index_values for vector)"
    )
    ci95: List[List[float]] = Field(
        description="95% credible intervals [[lo, hi], ...]"
    )
    units: str
    sample_size: Union[int, List[int]]
    sample_size_rationale: str

    # For vector-valued data (time courses, dose responses)
    index_values: Optional[List[float]] = None
    index_unit: Optional[str] = None
    index_type: Optional[IndexType] = None

    # Inputs and assumptions
    inputs: List[EstimateInput] = Field(
        description="Literature values with provenance"
    )
    assumptions: Optional[List[ModelingAssumption]] = None

    # Monte Carlo code
    distribution_code: str = Field(
        description="Python: derive_distribution(inputs, ureg) -> "
                    "{'median': [...], 'ci95': [...], 'units': str}"
    )
```

##### 5.9 Scenario and Intervention

For studies involving treatment, the `Scenario` captures the clinical context.

```
class Intervention(BaseModel):
    """A single treatment intervention."""
    agent: str = Field(description="Drug or treatment name")
    dose: Optional[str] = None
    route: Optional[str] = None
    schedule: Optional[str] = None

class Scenario(BaseModel):
    """Treatment scenario for intervention studies."""
    interventions: List[Intervention]
    treatment_line: Optional[str] = None
    prior_treatments: Optional[List[str]] = None
    description: str = Field(
```

```
        description="Narrative description of the clinical scenario"
    )
```

### 6 SubmodelTarget Schema Details

This section provides detailed field descriptions and representative YAML examples for each layer of the SubmodelTarget schema. These examples are referenced from the Methods section; the complete schema is defined in Section 5.

#### 6.1 Input Fields

Each input captures a single value extracted from literature with full provenance. The key fields are:

- `name`: Unique identifier referenced elsewhere in the schema
- `value` and `units`: The extracted numeric value with units
- `uncertainty`: Standard deviation or 95% confidence interval, when reported
- `n`: Sample size, for propagating uncertainty through the likelihood
- `input_type`: Classification as `direct_measurement`, `proxy_measurement`, `inferred_estimate`, or `assumed_value`
- `role`: How the input is used: `target` (likelihood term), `auxiliary` (supporting), `initial_condition`, or `fixed_parameter`
- `source_ref`: Reference to a literature source with DOI
- `value_snippet`: Quoted text from the source containing the value

Code block 1 shows a representative input.

```yaml
inputs:
  - name: pdgf_fold_mean
    value: 4.37
    units: dimensionless
    uncertainty:
      sd: 0.49
    n: 3
    input_type: direct_measurement
    role: target
    source_ref: Schneider2001
    source_location: Abstract
    value_snippet: "Cell proliferation (4.37 +/- 0.49-
                    and 2.96 +/- 0.39-fold of control)."
```

Figure 1: An input capturing a proliferation measurement with provenance.

#### 6.2 Parameter and Prior Specification

Each parameter includes a prior distribution with documented rationale. The schema supports four distribution families: LogNormal (positive parameters with multiplicative uncertainty), Normal (unconstrained), Uniform (bounded with no preference), and HalfNormal (positive with mode at zero).

Code block 2 shows a parameter specification.

```
calibration:
  parameters:
    - name: k_apsc_prolif
      units: 1/day
      prior:
        distribution: lognormal
        mu: 0.0      # log(1.0)
        sigma: 1.0
        rationale: |
          Wide prior centered at 1/day. A 4-fold increase
          in 1 day implies k ~ ln(4) ~ 1.4/day.
```

Figure 2: A parameter with lognormal prior and documented rationale.

#### 6.3 Forward Model Specification

The `forward_model` field captures the dynamics relevant to the experiment. For time-course data, this is typically a simplified ODE; for steady-state or derived measurements, an algebraic model may suffice. State variables and the independent-variable block are nested inside `forward_model`.

Code block 3 shows a forward model using the exponential growth template.

```
calibration:
  forward_model:
    type: exponential_growth
    rate_constant: k_apsc_prolif
    independent_variable:
      name: time
      units: day
      span: [0.0, 1.0]
    state_variables:
      - name: aPSC_rel
        units: dimensionless
        initial_condition:
          value: 1.0
          rationale: Normalized to control baseline.
    data_rationale: |
      BrdU assay measures proliferation over ~24h with
      constant PDGF stimulus. Exponential growth fits.
    submodel_rationale: |
      At saturating PDGF, the full model reduces to
      dN/dt = k_apsc_prolif * N.
```

Figure 3: Forward model specification with state variable and time span.

##### 6.4 Error Model Specification

The `error_model` field is a list of measurement entries, each specifying the statistical relationship between model predictions and one observed quantity. The `observation_code` derives the observed value and uncertainty from raw inputs. The `observable` defines the transformation from state variables to predicted measurement. The `likelihood` specifies the distribution family.

Code block 4 shows a complete error model entry.

```
error_model:
  - name: viability_48h
    units: dimensionless
    uses_inputs: [viability_mean, viability_sem]
    evaluation_points: [2.0]  # days
    sample_size: 3
    observable:
      type: identity
      state_variables: [aPSC_norm]
    observation_code: |
      def derive_observation(inputs, sample_size):
          mean = inputs['viability_mean']
          sem = inputs['viability_sem']
          sd = sem * np.sqrt(sample_size)  # SEM to SD
          return {'value': mean, 'sd': sd}
    likelihood:
      distribution: normal
      rationale: |
        Viability near 1 with small SD; normal adequate.
```

Figure 4: Error model specification linking forward model predictions to observed data.

### 7 Complete CalibrationTarget Example

This section presents a complete CalibrationTarget for baseline CD8+ T cell density in PDAC, demonstrating how full-model observables, reference database constants, and Monte Carlo distribution derivation work together.

#### 7.1 Overview

This target constrains baseline intratumoral CD8+ T cell density using immunohistochemistry data from Jansen et al. (2021), who measured CD8 density across 599 pancreatic cancers using tissue microarrays, of whom 444 had both CD8 IHC and DOG1 staging information and contribute to the mixture-of-lognormals distribution used here. The observable computes CD8+ density from full model species (effector and exhausted CD8+ T cells) and reference database constants (cancer cell geometry). The distribution code combines two patient subgroups via a mixture of lognormals.

#### 7.2 Observable

The observable maps full model species to the experimental measurement:

```
observable:
  code: |
    def compute_observable(time, species_dict, constants, ureg):
        import numpy as np
        cd8_eff = species_dict['V_T.CD8']
        cd8_exh = species_dict['V_T.CD8_exh']
        C1 = species_dict['V_T.C1']
```

```
        cross_section = constants['pdac_cancer_cell_cross_section']
        cellularity = constants['pdac_cellularity_fraction']

        total_cd8 = cd8_eff + cd8_exh
        tumor_area = C1 * cross_section / cellularity
        density = total_cd8 / tumor_area
        return density * ureg('cell / millimeter**2')
  units: cell / millimeter**2
  species: [V_T.CD8, V_T.CD8_exh, V_T.C1]
  constants:
    - name: pdac_cancer_cell_cross_section
      value: 0.000227
      units: millimeter**2 / cell
      biological_basis: |
        Mean cross-sectional area of PDAC cancer cells from
        histomorphometric measurements. Derived from mean cell
        diameter of 17 micrometers.
      source_type: reference_db
      reference_db_name: pdac_cancer_cell_cross_section
    - name: pdac_cellularity_fraction
      value: 0.15
      units: dimensionless
      biological_basis: |
        Fraction of tumor volume occupied by cancer cells in PDAC.
        PDAC tumors are characteristically desmoplastic with low
        cellularity (10-20%).
      source_type: reference_db
      reference_db_name: pdac_cellularity_fraction
  support: positive
  mapping_rationale: |
    IHC measures CD8+ cells per mm^2 of tissue section. The model
    tracks CD8+ T cells as counts in the tumor compartment (V_T).
    To convert model cell counts to density, we divide by tumor
    area estimated from cancer cell count, cross-section, and
    cellularity fraction.
  experimental_denominator: "mm^2 of tumor tissue section area"
  model_denominator_species: [V_T.C1]
```

##### 7.3 Empirical Data

The distribution code combines two patient subgroups (DOG1-negative and DOG1-positive) via mixture-of-lognormals Monte Carlo:

```
empirical_data:
  median: [138.8]
  ci95: [[20.3, 965.2]]
  units: cell / millimeter**2
  sample_size: 444
```

```
sample_size_rationale: |
  Total of 444 patients: 368 DOG1-negative + 76 DOG1-positive.
inputs:
  - name: mean_cd8_stroma_neg
    value: 227.7
    units: cell / millimeter**2
    source_ref: Jansen2021
    source_location: "Table 4"
    value_snippet: "CD8 ... 227.7 +/- 15.0"
  - name: sem_cd8_stroma_neg
    value: 15.0
    units: cell / millimeter**2
    source_ref: Jansen2021
    source_location: "Table 4"
    value_snippet: "CD8 ... 227.7 +/- 15.0"
  - name: n_stroma_neg
    value: 368
    units: dimensionless
    source_ref: Jansen2021
    source_location: "Table 1"
    value_snippet: "DOG1 negative 368 (61.4%)"
  - name: mean_cd8_stroma_pos
    value: 220.1
    units: cell / millimeter**2
    source_ref: Jansen2021
    source_location: "Table 4"
    value_snippet: "CD8 ... 220.1 +/- 33.0"
  - name: sem_cd8_stroma_pos
    value: 33.0
    units: cell / millimeter**2
    source_ref: Jansen2021
    source_location: "Table 4"
    value_snippet: "CD8 ... 220.1 +/- 33.0"
  - name: n_stroma_pos
    value: 76
    units: dimensionless
    source_ref: Jansen2021
    source_location: "Table 1"
    value_snippet: "DOG1 positive 76 (12.7%)"
distribution_code: |
  def derive_distribution(inputs, ureg):
      import numpy as np
      rng = np.random.default_rng(42)
      n_mc = 100000

      # Subgroup 1: DOG1-negative
      mean1 = inputs['mean_cd8_stroma_neg'].magnitude
      sem1 = inputs['sem_cd8_stroma_neg'].magnitude
```

```
        n1 = int(inputs['n_stroma_neg'].magnitude)
        sd1 = sem1 * np.sqrt(n1)

        # Subgroup 2: DOG1-positive
        mean2 = inputs['mean_cd8_stroma_pos'].magnitude
        sem2 = inputs['sem_cd8_stroma_pos'].magnitude
        n2 = int(inputs['n_stroma_pos'].magnitude)
        sd2 = sem2 * np.sqrt(n2)

        # Convert to lognormal parameters
        def to_lognormal(m, s):
            var = s**2
            mu = np.log(m**2 / np.sqrt(m**2 + var))
            sigma = np.sqrt(np.log(1 + var / m**2))
            return mu, sigma

        mu1, sig1 = to_lognormal(mean1, sd1)
        mu2, sig2 = to_lognormal(mean2, sd2)

        # Mixture sampling
        w1 = n1 / (n1 + n2)
        mask = rng.random(n_mc) < w1
        samples = np.where(
            mask,
            rng.lognormal(mu1, sig1, n_mc),
            rng.lognormal(mu2, sig2, n_mc)
        )

        return {
            'median': [float(np.median(samples))],
            'ci95': [[float(np.percentile(samples, 2.5)),
                      float(np.percentile(samples, 97.5))]],
            'units': 'cell / millimeter**2'
        }
```

##### 7.4 Source and Context

```
primary_data_source:
  doi: "10.7717/peerj.11397"
  title: "Expression of immune cell markers in the tumor
          microenvironment and association with survival"
  authors: [Jansen, van Doorn, et al]
  year: 2021
  source_tag: Jansen2021

experimental_context:
  species: human
  system: clinical_resection
```

```
  indication: PDAC
  treatment_status: treatment_naive
  stage: resectable

study_interpretation: |
  Jansen et al. measured CD8+ T cell density across 599 pancreatic
  cancers using tissue microarrays and digital image analysis. We
  combine two subgroups (DOG1-positive and DOG1-negative) as a
  mixture of lognormals to capture the full population distribution.

key_assumptions:
  - "CD8 IHC detects both effector and exhausted T cells equally"
  - "Reported +/- values are SEM (confirmed by sqrt(n) test)"
  - "DOG1 expression does not significantly affect CD8 density"

key_study_limitations:
  - "TMA cores (0.6mm) may not represent whole-slide heterogeneity"
  - "Mixed pathologic stages without neoadjuvant therapy reporting"
  - "2D histology section vs. 3D model compartment"
```

### 8 SubmodelTarget Source Characteristics

This section provides detailed breakdowns of source quality, species translation, and indication match for the 37 curated SubmodelTarget extractions.

#### 8.1 Source Quality

Source quality categorizes the type of experimental evidence. *Human clinical* denotes data from human patient samples or clinical measurements. *Human in vitro* denotes experiments using human-derived cell lines or primary human cells in culture. *Animal in vivo* denotes measurements from animal models (e.g., mouse xenografts). *Animal in vitro* denotes experiments using animal-derived cells in culture (e.g., murine splenocyte assays, rat stellate cell cultures).

Table 2: Distribution of primary data source quality across SubmodelTargets.

| Source Quality | Count | % |
|---|---|---|
| primary_human_clinical | 17 | 46% |
| primary_animal_in_vivo | 9 | 24% |
| primary_animal_in_vitro | 7 | 19% |
| primary_human_in_vitro | 3 | 8% |
| review_article | 1 | 3% |
| **Total** | **37** | |

#### 8.2 Species Translation

Species translation describes the mapping from the experimental organism to the model target species (human). Targets labeled human→human required no cross-species translation. Mixed-species entries (e.g., human and rat→human) indicate that the extraction combined data from multiple species, such as a human clinical measurement anchored by rat-derived kinetic parameters.

Table 3: Species translation from data source to model target. Human→human indicates no cross-species translation required.

| Translation | Count | % |
|---|---|---|
| human→human | 21 | 57% |
| mouse→human | 13 | 35% |
| rat→human | 3 | 8% |
| **Total** | **37** | |

##### 8.3 Indication Match

Indication match describes how closely the experimental disease context matches the PDAC model target. *Exact*: data from PDAC patients or PDAC-derived models. *Proxy*: data from a different indication used as a mechanistic proxy (e.g., other cancer types, chronic pancreatitis, fibrosis models with similar stromal biology). *Related*: data from the same organ or disease class but not a direct match.

Table 4: How closely the data source indication matches the model target (PDAC).

| Indication Match | Count | % |
|---|---|---|
| Data from PDAC patients/models | 17 | 46% |
| Proxy indication (e.g., other cancers, fibrosis) | 16 | 43% |
| Related indication (same organ/disease class) | 4 | 11% |
| **Total** | **37** | |

#### 9 CalibrationTarget Detailed Metrics

This section provides detailed metrics for the 45 CalibrationTargets applied to the PDAC QSP model.

##### 9.1 Observable Categories

##### 9.2 Observable Complexity

The most complex observables (19 species) are neoadjuvant immunotherapy targets computing immune subset percentages of all nucleated cells, requiring all modeled immune, stromal, and tumor populations in the denominator. The highest-constant observables (6 constants) require geometric and cellularity reference parameters for converting between model cell counts and histologic measurements. Across all targets, observables reference 25 unique model species spanning immune cells ($CD8^+$ effector and exhausted, Treg, Th, NK), myeloid cells (M1/M2 macrophages, MDSC, cDC1/cDC2), stromal cells (myCAF, iCAF, apCAF, quiescent PSC), tumor cells, ECM, and microenvironment variables (VEGF, lactate, pO2, glucose, HIF).

Table 5: CalibrationTarget observable categories.

| Observable Category | Count |
|---|---|
| Immune/stromal fractions | 25 |
| Cell densities | 8 |
| Population ratios | 7 |
| Fold-changes | 2 |
| Clinical endpoints | 2 |
| Metabolic/TME markers | 1 |

Table 6: Observable complexity across 45 CalibrationTargets.

| Metric | Mean | Min | Max |
|---|---|---|---|
| Model species per observable | 8.4 | 1 | 19 |
| Named constants per observable | 0.7 | 0 | 6 |
| Empirical inputs per target | 6.4 | 1 | 18 |

#### 9.3 Extraction Model Attribution

Table 7: CalibrationTarget extraction model breakdown.

| Extraction Model | Count | Percentage |
|---|---|---|
| Claude Opus 4.6 | 27 | 60% |
| GPT-5.1 | 18 | 40% |
| **Total** | **45** | |

Of the 45 AI-generated targets, 30 (66%) are tagged as human-verified, indicating that a domain expert reviewed the complete target after initial extraction. 15 targets (33%) include data digitized from figures, where per-patient data points were manually extracted and the CalibrationTarget structure was assembled by the LLM from those data points.

#### 9.4 Primary Data Sources

The 45 targets drew from 18 unique primary sources (by DOI), with publication years spanning 2007–2026. Two sources contributed the majority of targets: a neoadjuvant immunotherapy trial (Li et al., Cancer Cell, 2022) provided data for all 18 GVAX/nivolumab targets, and a large-scale immunohistochemistry study (Liudahl et al., Cancer Discovery, 2021) provided data for 10 baseline targets covering immune cell densities, fractions, and ratios.

### 10 Inference Results

All 37 SubmodelTargets were translated to a joint Julia inference script using Turing.jl[1]. The translator identified 19 unique parameters and generated corresponding priors and likelihood terms. The plurality of targets (49%) use closed-form algebraic forward models; the remainder use structured model types including steady-state, batch accumulation, and first-order decay models (Table 8).

Table 8: Distribution of forward model types used in calibration submodels.

| Model Type | Count | % |
|---|---|---|
| Algebraic (closed-form) | 18 | 49% |
| Steady-state ratio | 5 | 14% |
| Batch accumulation | 4 | 11% |
| Steady-state density | 4 | 11% |
| Steady-state fraction | 3 | 8% |
| First-order decay ODE | 2 | 5% |
| Steady-state concentration | 1 | 3% |
| **Total** | **37** | |

The generated script is self-contained, requiring only DifferentialEquations.jl, Turing.jl, and Distributions.jl[2]. Each target contributes a simulate function, observed data constants, and a likelihood term. In this application, most targets constrain individual parameters through algebraic or structured forward models (steady-state, batch accumulation, first-order decay), with 11 parameters shared across multiple targets. The framework supports arbitrarily complex scenarios: multiple derivations constraining the same parameter, single targets constraining multiple parameters, and forward models involving ODEs that cannot be analytically inverted.

##### 10.1 Convergence Diagnostics

We ran 4 chains of 1000 samples each using the NUTS sampler. All parameters achieved $\hat{R} < 1.01$, with bulk-ESS $> 2600$ and tail-ESS $> 1300$ throughout (Table 9). Sampling took about 42 seconds on an M-series MacBook Air.

##### 10.2 Posterior Distributions

Figure 5 shows posterior estimates for all 19 parameters. Table 10 reports medians and 90% credible intervals.

Most posteriors are narrower than their priors, indicating that the data constrained the parameters. The tumor growth rate `k_C1_growth` (5.34e-03 day$^{-1}$, 90% CI: 2.22e-03–0.0128) implies a doubling time of about 130 days, consistent with clinical PDAC volume-doubling-time measurements[3,4]. The activated PSC death rate `k_apsc_death` (0.0388 day$^{-1}$) implies a half-life of about 18 days.

The IL-2-dependent division counts for CD4+ T cells (8.03) fall within the 7–8 generation range expected from CFSE proliferation tracking, where dye dilution limits counting. The CD8+ estimate (5.31) falls below this range, suggesting either a lower IL-2 division capacity for the CD8+ subset in the source data or unmodelled differences between the CD4 and CD8 experimental conditions. Only one calibration target constrains this parameter, its posterior concentrates near the lower end of the prior (Table 10) rather than spreading widely, and it shows the lowest tail-ESS in the corpus (Table 9), so the estimate should be interpreted with caution.

#### 11 Model-Aware Prompt Construction Details

For each target, the prompt injects mechanistic context extracted from the model structure. For target parameters, the system automatically retrieves: the parameter's name, units, and biological

Table 9: Convergence diagnostics for MCMC sampling. All $\hat{R} < 1.01$ indicates convergence.

| Parameter | $\hat{R}$ | ESS (bulk) | ESS (tail) |
|---|---|---|---|
| N_IL2_CD4 | 1.000 | 4598 | 2549 |
| N_IL2_CD8 | 1.002 | 2846 | 1334 |
| cd8_exclusion_fraction | 1.001 | 2705 | 2325 |
| k_C1_growth | 1.001 | 4912 | 3195 |
| k_CCL2_deg | 1.001 | 5256 | 2763 |
| k_CCL2_sec | 1.001 | 5453 | 2998 |
| k_CD8_death | 1.001 | 2851 | 2889 |
| k_M1_pol | 1.001 | 3020 | 2396 |
| k_M2_pol | 1.001 | 2773 | 2542 |
| k_MDSC_death | 1.001 | 2679 | 2515 |
| k_MDSC_rec | 1.001 | 2657 | 2621 |
| k_Mac_death | 1.000 | 3154 | 2987 |
| k_Mac_rec | 1.001 | 3055 | 2861 |
| k_Treg_death | 1.000 | 3813 | 2318 |
| k_apsc_death | 1.000 | 4222 | 2208 |
| k_psc_activation | 1.000 | 4511 | 3033 |
| n_CD8_clones | 1.001 | 4657 | 2895 |
| q_CD8_T_in | 1.000 | 4492 | 2440 |
| q_Treg_T_in | 1.002 | 5077 | 2879 |

Table 10: Posterior parameter estimates from joint Bayesian inference. Credible intervals are 90% highest density intervals. †Auxiliary parameter introduced during extraction.

| Parameter | Median | 90% CI | Units |
|---|---|---|---|
| N_IL2_CD4 | 8.03 | [5.73, 10.3] | dimensionless |
| N_IL2_CD8 | 5.31 | [4.03, 5.99] | dimensionless |
| cd8_exclusion_fraction† | 0.912 | [0.741, 0.982] | dimensionless |
| k_C1_growth | 5.34e-03 | [2.22e-03, 0.0128] | $\text{day}^{-1}$ |
| k_CCL2_deg | 4.29 | [2.77, 6.75] | $\text{hour}^{-1}$ |
| k_CCL2_sec | 3.32e-10 | [2.57e-10, 4.33e-10] | $\text{nmol}\cdot\text{cell}^{-1}\cdot\text{day}^{-1}$ |
| k_CD8_death | 2.23 | [0.546, 7.55] | $\text{day}^{-1}$ |
| k_M1_pol | 0.0392 | [0.0298, 0.0519] | $\text{day}^{-1}$ |
| k_M2_pol | 0.0776 | [0.0545, 0.111] | $\text{day}^{-1}$ |
| k_MDSC_death | 0.08 | [0.0568, 0.114] | $\text{day}^{-1}$ |
| k_MDSC_rec | 1.36e+07 | [9.76e+06, 1.92e+07] | $\text{cell}\cdot\text{mL}^{-1}\cdot\text{day}^{-1}$ |
| k_Mac_death | 0.0219 | [5.42e-03, 0.0812] | $\text{day}^{-1}$ |
| k_Mac_rec | 8.92e+05 | [2.11e+05, 4.08e+06] | $\text{cell}\cdot\text{mL}^{-1}\cdot\text{day}^{-1}$ |
| k_Treg_death | 0.2 | [0.0664, 0.636] | $\text{day}^{-1}$ |
| k_apsc_death | 0.0388 | [0.0278, 0.0506] | $\text{day}^{-1}$ |
| k_psc_activation | 0.5 | [0.342, 0.737] | $\text{day}^{-1}$ |
| n_CD8_clones | 10.3 | [8.66, 12.2] | dimensionless |
| q_CD8_T_in | 4.03e-04 | [3.52e-04, 4.60e-04] | $\text{min}^{-1}\cdot\text{cm}^{-3}$ |
| q_Treg_T_in | 6.47e-04 | [4.63e-04, 9.05e-04] | $\text{min}^{-1}\cdot\text{cm}^{-3}$ |

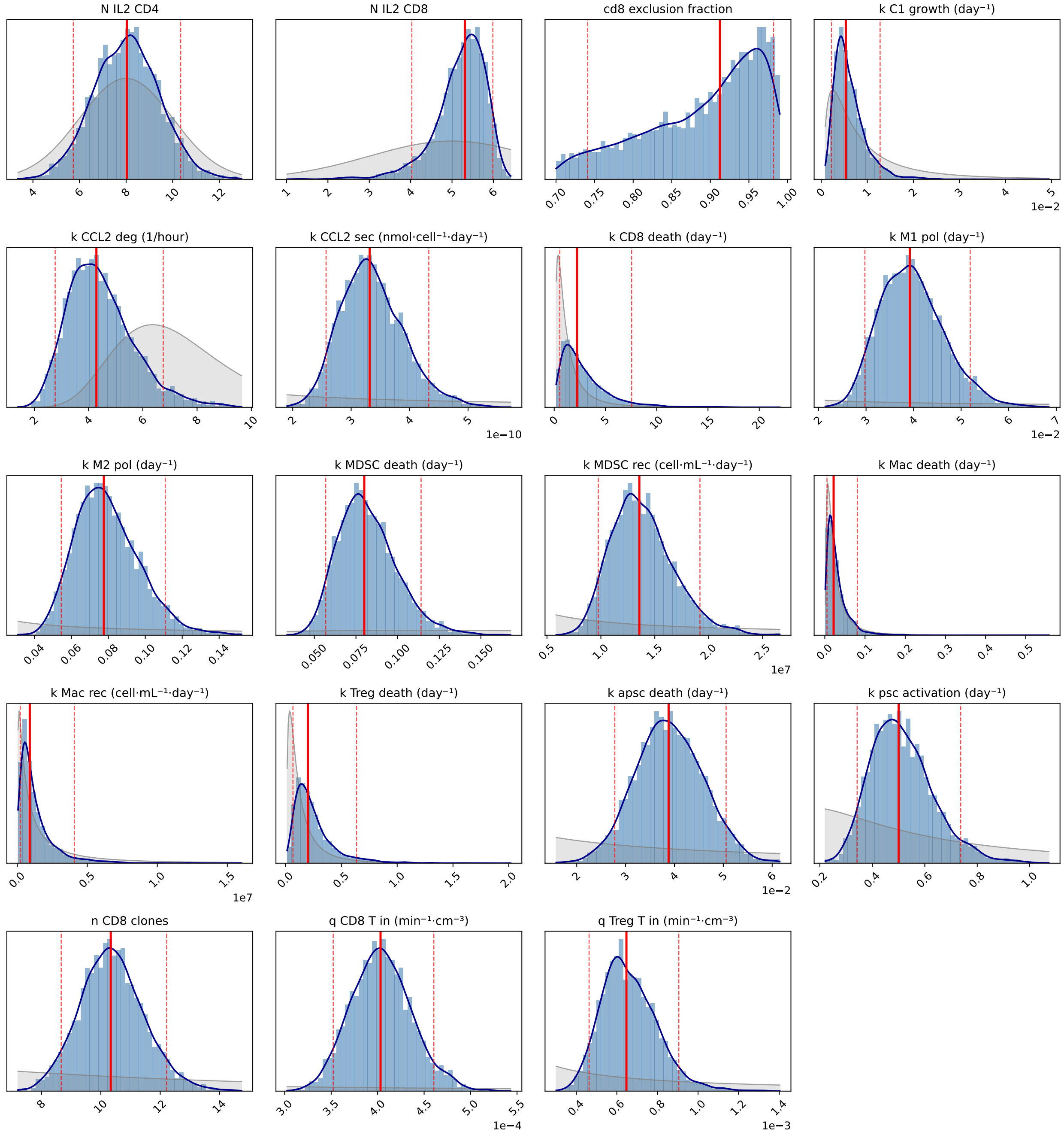

Figure 5: Posterior distributions from joint Bayesian inference. Gray shading shows the prior distribution; blue histogram and line show the posterior. Red lines indicate median (solid) and 90% credible interval bounds (dashed).

description; the reactions where the parameter appears with their rate laws; the species involved in those reactions with descriptions; other parameters appearing in the same reactions; and the broader reaction network showing other reactions that involve the same species. For target observables, the system retrieves the observable's name, units, and description, along with the species and compartments that define it. For example, when searching for data to calibrate a proliferation rate `k_apsc_prolif`, the prompt includes the reaction `aPSC -> 2*aPSC` with rate law `k_apsc_prolif * aPSC`, species descriptions explaining that aPSC represents activated pancreatic stellate cells, and related parameters like death rates that appear in the same reaction network.

This context guides the LLM's search toward experiments that actually constrain the targets, rather than experiments that mention similar concepts but measure different quantities. For parameters, the broader reaction network helps the LLM identify whether additional parameters should be jointly calibrated. Extracting multiple parameters from the same study is intentional and preferred when the data supports it, as shared experimental conditions reduce inter-study variability.

To encourage literature diversity, source exclusion prevents the LLM from reusing papers across extractions. The prompt includes a list of primary sources already used for other targets, and a strict exclusion mode extends this to sources used by related parameters in the same reaction network.

Beyond mechanistic context, the prompt includes the complete schema structure with field descriptions, enabling the LLM to generate valid output without trial and error. The prompt also provides explicit guidance on common pitfalls: distinguishing SEM from SD (with conversion formulas), avoiding hardcoded numeric values in code blocks, and properly documenting source relevance for cross-species or cross-indication extractions. By showing validation requirements upfront (such as the prohibition on non-peer-reviewed sources and required uncertainty thresholds for proxy data), the prompt reduces the retry loop by preventing predictable errors.

### 12 Schema Implementation Details

Both schemas are implemented using Pydantic[5], a Python library for declarative data validation. Each schema component is defined as a Pydantic model with typed fields and constraints. When YAML is parsed, Pydantic validates all constraints automatically: required fields must be present, values must match declared types, enumerations must use allowed values, and cross-field constraints must be satisfied.

The validation error messages are detailed and structured, reporting the field path, expected type, and actual value for each violation. This specificity is important for the LLM retry loop: when extraction fails validation, the error message provides actionable feedback. For example, a snippet validation error reports "Value-snippet mismatches (possible hallucination): input 'cell_count_mean' has value 4.37 but snippet contains '3.21'. Check that extracted values match the source text exactly." This tells the LLM both what failed and how to fix it.

SubmodelTarget uses discriminated unions for its 15 forward model types: the `type` field determines which additional fields are required, ensuring type-specific validation without manual dispatch logic. CalibrationTarget uses a class hierarchy with shared base models for source documentation, relevance assessment, and experimental context.

### 13 Source Relevance Assessment Details

Each target includes structured metadata about its data sources. The `primary_data_source` block captures the main reference: DOI, title, authors, year, and a short `source_tag` used for crossreferencing elsewhere in the schema. Secondary sources that provide supporting context (e.g., reviews explaining biological mechanisms, or related studies used for uncertainty estimation) are listed in `secondary_data_sources`, each with a `contribution` field explaining how that source informs the extraction.

The `indication_match` field classifies how well the source disease context matches the model: `exact` (same indication), `related` (same organ or disease class), `proxy` (different tissue used as mechanistic proxy), or `unrelated`. Cross-indication extractions require a justification explaining why the data is still informative.

Species translation is captured by `species_source` and `species_target` fields. When these differ (e.g., mouse data informing a human model), the extraction must document the assumed translation.

The `source_quality` field categorizes the evidence type: `primary_human_clinical`, `primary_human_in_vitro`, `primary_animal_in_vivo`, `primary_animal_in_vitro`, `review_article`, or `textbook`. By default, non-peer-reviewed sources trigger a validation warning; in practice, preprints (e.g., from bioRxiv) that have been publicly available for several months can provide valuable data, and the validator can be configured to accept them with documented justification.

For immune and stromal parameters, `tme_compatibility` assesses whether the tumor microenvironment in the source matches the target model (`high`, `moderate`, or `low`). A T cell trafficking rate measured in melanoma (a T cell-permissive tumor) may not translate directly to PDAC (an immunosuppressive tumor); low compatibility requires explicit documentation.

Finally, `estimated_translation_uncertainty_fold` quantifies the expected uncertainty from all translation factors combined. Cross-species extraction from a proxy indication with low TME compatibility might warrant 10-fold or greater uncertainty, which propagates to prior width during inference.

### 14 Collaboration Mode Details

The two collaboration modes observed in this study offer different trade-offs. In the batch pipeline mode, the LLM produces a structured draft that the modeler then curates interactively: selecting appropriate forward model types and prior distributions for SubmodelTargets, correcting observable code and species mappings for CalibrationTargets, and revising source relevance assessments for both. This mode is efficient for initial coverage (many files per run) but carries a high rejection rate (58% in a documented batch review) because the LLM lacks real-time modeler guidance on source selection, species mapping, and denominator definitions. In the interactive mode, the modeler and LLM work together from the start, with the modeler directing extraction decisions while the LLM handles text generation, code writing, and literature comprehension. This mode front-loads modeler effort during extraction and leaves less to revise afterwards. Because the batch and interactive runs used different LLMs (GPT-5.1 and Claude Opus 4.6) and the interactive work followed curation of the batch outputs, this difference is observational rather than evidence that interactive extraction is inherently superior.

In both modes, the schema serves as the collaboration interface. It defines what information must be captured (ensuring completeness), constrains how it is represented (enabling validation), and separates data extraction from modeling decisions (clarifying where different types of expertise are needed). The validators provide guardrails during both LLM extraction and human editing, catching errors regardless of their source.

#### 14.1 Categories of modeler-supplied context

The field-level curation rates reported in Section 3.2 of the main text aggregate changes of several different kinds. Across the SubmodelTarget and CalibrationTarget revisions, modeler-supplied context fell into the following recurring categories:

1. *Unit conventions and dimensional bridges.* Conversion factors that the LLM cannot infer from `model_structure.json` alone, such as a missing $10^{-6}$ tissue protein density conversion in cDC and Treg density formulas, or the Abercrombie 3D-to-2D correction used to convert section-level cell counts to volumetric densities.

2. *Source selection and retirement.* Excluding a study after recognizing a mismatch between assay or marker definitions and model categorization (e.g., a study reporting CD86+ as an M1 marker, where the simulation does not enforce that definition), or substituting a better source when one becomes available.

3. *Cross-source aggregation.* Decisions to pool measurements from multiple studies (e.g., `cd8_density` aggregated across Golesworthy 2022 and Liu 2015) rather than treat them as separate targets.

4. *Measurement-to-state translation logic.* Code inside the observable function that converts a paper's measurement into a model-relevant quantity, such as serum-to-tissue bridging factors that translate plasma cytokine concentrations to tissue-level model species, or active-fraction adjustments where immunoassays detect total cytokine although only a signalling-active fraction is biologically meaningful.

5. *Denominator and normalization choices.* What a percentage is taken of (CD8 of CD3 versus of nucleated cells), and which compartment volume normalizes a per-tissue measurement.

6. *Scenario-specific forward-model structure.* Constraints particular to a calibration scenario rather than to the underlying biology: peak-ordering constraints (e.g., a tumor-volume minimum that must follow a CD8 peak in the GVAX-only arm), or reparameterization of a kinetic in its linear sub-saturating regime when the data do not span saturation.

7. *Uncertainty widening from source quality.* Adjusting prior or likelihood widths upward when a source is cross-species or cross-indication; the schema's `source_relevance` block records the rationale alongside the widening.

These categories did not contribute equally to curation effort. Unit conventions and source selection accounted for the largest share of edits. Auxiliary-parameter additions and denominator decisions occurred less often but represented modeling decisions about how a measurement maps to a model state, and typically took longer per edit than the unit corrections. Forward-model restructuring and uncertainty widening were applied selectively to a smaller number of targets.

### 15 Detailed Comparison to Existing Approaches

General-purpose biomedical extraction frameworks like LLM-IE[6] provide building blocks for named entity recognition and relation extraction, but do not address the downstream requirements of QSP calibration: uncertainty quantification, forward model specification, and code generation for inference. Interactive systems like SciDaSynth[7] support structured data extraction with human-in-the-loop refinement, but focus on tabular output rather than the hierarchical schema needed for QSP model calibration. QSP-Copilot[8] applies LLMs to QSP model development broadly, including literature extraction, but emphasizes model structure discovery (entity-interaction pairs) rather than quantitative parameter calibration with provenance.

The validation approach in MAPLE draws on established strategies for hallucination detection. Requiring extracted values to appear in quoted source text is analogous to the grounding strategies used in retrieval-augmented generation[9], where LLM outputs are verified against retrieved documents. The schema-based validation enforces structural constraints similar to those in constrained decoding and structured output systems, where JSON schemas guide LLM generation toward valid output[10]. The key contribution is combining these strategies with domain-specific validators (DOI resolution, unit parsing, code execution) in a framework tailored to quantitative scientific extraction.

Recent work on knowledge graph construction from biomedical literature[11,12] demonstrates that LLMs can extract structured relationships at scale. The Immune Cell Knowledge Graph (ICKG) used zero-shot prompting with Llama 3.1 to extract activation and inhibition relationships from over 24,000 PubMed abstracts. Such approaches prioritize coverage and can tolerate some extraction errors because downstream analyses aggregate across many relationships. Parameter calibration has different requirements: each extracted value directly influences model predictions, so individual errors matter more. MAPLE trades some throughput for higher per-target validation stringency.

The IQ Consortium has outlined best practices for AI/ML in quantitative modeling, including the use of NLP for knowledge extraction from biomedical literature[13]. They emphasize explainability, human-in-the-loop oversight, and consideration of context of use. MAPLE aligns with these principles: the schema documents reasoning and provenance, validators catch errors before human review, and the generated code is transparent and modifiable. The growing interest in coupling QSP with machine learning[14,15,16,17] suggests that hybrid approaches combining mechanistic modeling with data-driven methods will become increasingly important; automated literature extraction is one component of this broader integration. Recent work has also explored using LLMs directly as tools for comparing and synthesizing perspectives on QSP methodology[18]. While we evaluate MAPLE on a QSP model, the approach is not limited to pharmacology: any parameter-rich mechanistic model that draws calibration data from literature (e.g., systems biology signaling networks, physiologically-based models) could benefit from structured extraction with validation.

[4] Jang KM, Kim SH, Kim YK, Song KD, Lee SJ, Choi D. Missed Pancreatic Ductal Adenocarci-

noma: Assessment of Early Imaging Findings on Prediagnostic Magnetic Resonance Imaging. European Journal of Radiology. 2015;84(8):1473-9. doi:10.1016/j.ejrad.2015.05.012.
